# Ancient DNA reveals historical demographic decline and genetic erosion in the Atlantic bluefin tuna

**DOI:** 10.1101/2024.09.14.613028

**Authors:** Adam Jon Andrews, Emma Falkeid Eriksen, Bastiaan Star, Kim Præbel, Antonio Di Natale, Estrella Malca, Glenn Zapfe, Vedat Onar, Veronica Aniceti, Gabriele Carenti, Gäel Piquès, Svein Vatsvåg Nielsen, Per Persson, Federica Piattoni, Francesco Fontani, Lane M. Atmore, Oliver Kersten, Fausto Tinti, Elisabetta Cilli, Alessia Cariani

**Author notes:** EC and AC should be considered joint senior author.

## Abstract

Overexploitation has depleted fish stocks during the past century, nonetheless its genomic consequences remain poorly understood. Characterising the spatiotemporal patterns of these consequences may provide baseline estimates of past diversity and productivity to aid management targets, help predict future dynamics, and facilitate the identification of evolutionary factors limiting fish population recovery. Here, we evaluate human impacts on the evolution of the iconic Atlantic bluefin tuna (*Thunnus thynnus*), one of the longest and most intensely exploited marine fishes, with a tremendous cultural and economic importance. We sequenced whole genomes from modern (n=49) and ancient (n=41) specimens dating up to 5000 years ago, uncovering several novel findings. First, we identify temporally stable patterns of population admixture, as bluefin tuna caught off Norway and in the eastern Mediterranean share a greater degree of ancestry with Gulf of Mexico bluefin tuna than western and central Mediterranean bluefin tuna. This suggests that Atlantic spawning areas are important mixing grounds for the genetic diversity of Mediterranean bluefin tuna. We model effective population size to show that Mediterranean bluefin tuna began to undergo a demographic decline by the year 1900 to an extent not observed across the previous millennia. Coinciding with this, we found that heterozygosity and nucleotide diversity was significantly lower in modern (2013-2020), than ancient (pre-1941) Mediterranean bluefin tuna, suggesting bluefin tuna underwent a genetic bottleneck. With this work we show how ancient DNA provides novel perspectives on ecological complexity with the potential to inform the management and conservation of fishes.

**Significance:** Achieving the aim of the current UN Ocean Decade to “protect and restore ecosystems and biodiversity” is stymied by a lack of historical knowledge on how human exploitation has impacted and therefore what should be restored. Here, we sequence DNA in ancient fish bones to evaluate the historical diversity of the Atlantic bluefin tuna; which has been of great commercial importance for centuries. We find that bluefin tuna began to undergo demographic decline by 1900, 70 years earlier than currently recognised. Correspondingly, we find modern bluefin tuna had lower levels of genetic diversity than historical ones. This suggests that human impacts on the diversity of marine fishes are likely to have begun earlier and be more complex than previously thought.

## Main text

As we restore our oceans, fundamental questions remain as to how productive fish populations were historically and whether their recovery and adaptability will be hampered by genetic changes like a loss of genetic diversity, which have resulted from their exploitation (1, 2). Between 1970-2007, the ecologically, culturally and commercially important pelagic fish, Atlantic bluefin tuna (*Thunnus thynnus*) was severely depleted, resulting in the loss of large size-classes and rarity across its range; where it periodically disappeared from habitats such as the North, Norwegian and Black Seas (3–8). Fisheries data from recent decades estimate that between 1960-2009, the eastern stock abundance and range declined by an estimated 70% and 46-53%, respectively (4, 5). Mediterranean bluefin tuna stock abundance has recently recovered to 1970s levels following several years of international management and favourable oceanographic conditions, while it has returned to most of its previous habitats (9–13).

Bluefin tuna recovery is duly considered a fisheries management success (3, 14); as a result, catch quotas have increased since 2018 (13). However, historical sources suggest that the intensification of bluefin fisheries–and subsequent population declines–had begun centuries previous (15), indeed bluefin tuna have been commercially fished in the Mediterranean since before ancient Greek times. This trend has been shown for other marine populations and taxa (e.g., (16–19)), yet, little is known whether 1970 abundance baselines underestimate historical bluefin tuna productivity, and how centuries of exploitation have impacted its population structure, gene flow and genetic diversity. This information could be useful to predict future population dynamics, protect sustainability and maximise fisheries productivity (20–23). With this work, we use ancient DNA and whole genome sequencing to address these questions.

Bluefin tuna is characterised by its large size (up to 3.3 m in length and 725 kg in weight), far-reaching and inshore migration behaviour, and slow maturation (between 3-6 years, (24, 25)). Recent genomic studies (26, 27) support the delineation of two highly-connected populations, which are managed as two stocks (Figure 1). These are a western Atlantic component that spawns predominantly in the Gulf of Mexico (28) and an eastern Atlantic and Mediterranean component that spawns predominantly in several high-productivity areas of the Mediterranean Sea (25). Mature and juvenile individuals of both populations migrate into the Atlantic Ocean to feed,(29, 30), and exhibit high-levels of mixing (26, 27). The role of other understudied and potential spawning areas such as those in the Azores, Canary Islands and the Bay of Biscay (25, 31) are yet to be clearly defined, but bluefin tuna born in the Slope Sea (East of Cape Hatteras, USA, (28, 32)) appear to be a genetically mixed component of the two populations (27, 33). Due to the recent mismatch in recovery between the stocks and uncertainties on the scale of the decline in the western population(s) in the 1960s (3), there is concern about the genetic homogenization of the populations (33). In addition, there are many unknowns regarding migration and population structure, and while tagging and historical fishery data suggests that a portion of Mediterranean bluefin tuna is resident year-round (6, 34–36), there is as yet no genetic evidence for this.

**Figure 1.**
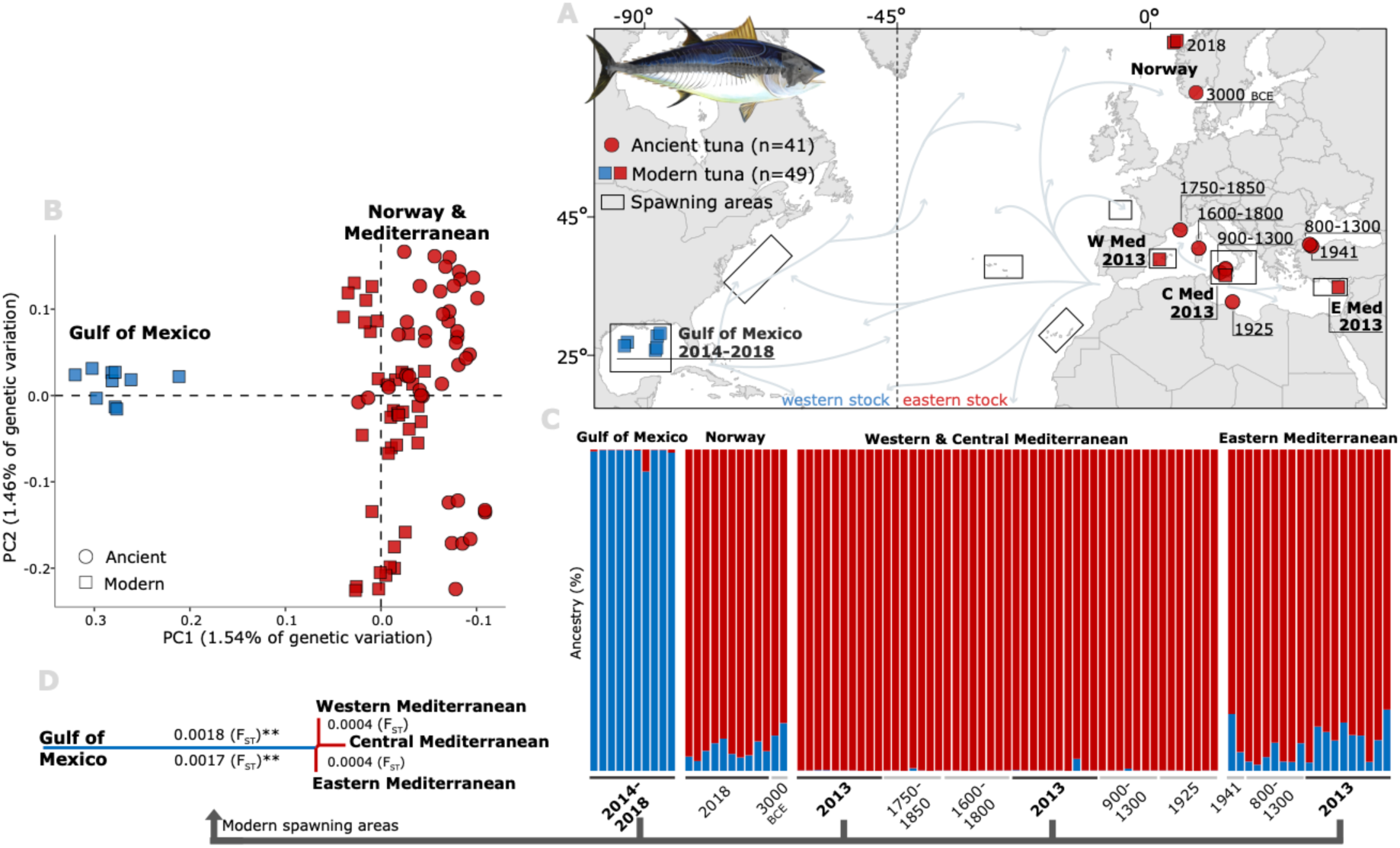
Spatial patterns in admixture of the two Atlantic bluefin tuna (*Thunnus thynnus*) populations consistent over millennia, and map (A) of samples sequenced, spanning over millennia across the species range. Samples (named by year) are coloured by stock: western stock (blue), eastern stock (red). Ancient samples analysed (n=41) are depicted using circles, modern (n=49) using squares. Modern larvae or YoY (young-of-year) samples from spawning sites are: Gulf of Mexico, Western (W Med), Central (C Med) and Eastern (E Med) Mediterranean. Spawning areas are approximately depicted using rectangles, including Atlantic sites not sampled. Grey arrows indicate stock distribution, mixing and migration routes. Stock management line at 45°W is illustrated as a dashed black line. Map created using ESRI ArcMap (v.10.6, https://arcgis.com). Principal components analysis (PCA, B) using 12,838 SNPs show two main sample clusters representing the western and eastern stock. STRUCTURE barplot (C) using the same dataset shows ancestry proportions for the optimal number of populations (K=2), tested using the Evanno Method, and evidence for increased admixture between stocks in Norwegian and Eastern Mediterranean individuals, consistent across millennia. Pairwise F_ST_ neighbour joining tree (D) shows differentiation between modern spawning area samples supports STRUCTURE results at 386,963 loci, indicating statistical significance using FDR-corrected *p*-values (**; p <0.001).

For millennia, tuna traps have iconically lined eastern Atlantic and Mediterranean coasts, intercepting spawning migrations of highly-revered BFT between April-September (15, 29). By the 1600-1800s, Mediterranean bluefin tuna shifted habitat and diet preferences further from shore (37) and catches fluctuated heavily (38) indicating that the population was dynamic and varied potentially due to climate or early anthropogenic impacts. From 1800-1900, large bluefin tuna were increasingly targeted in trap fisheries, indicating the potential for fisheries-induced evolution (39). Trap records suggest that by the 1880s, landings had reached comparable levels to the 1980s-2000s, the most intensive decades in BFT exploitation (3, 40). By 1970, the bluefin tuna range contraction away from the North, Norwegian and Black Seas and the closure of the majority of the trap fisheries–suggests that, despite the intense exploitation, the fishery’s productivity was depressed (6). Therefore, we hypothesised that longer-term data is required to investigate demographic changes, and its impact, prior to the 1970s, from when routine fisheries data exists.

Genomic analyses of temporal samples before and after overexploitation events provide key opportunities to test for the onset of overexploitation and its impact on population structure and genetic diversity (22, 41, 42). In studies on fishes, recent demographic history has been modelled using modern samples to reveal pre-industrial exploitation impacts in Atlantic herring (*Clupea harengus*) (16), but analyses directly testing genetic changes between temporally samples – e.g., to reveal signatures of genetic erosion – remains rare. This approach has been demonstrated in taxa like seabirds (43), (23, 42). Genetic erosion (the loss of genetic diversity e.g. heterozygosity, nucleotide diversity) may leave populations with lowered adaptive capacity, and remains poorly studied in marine fishes due to technical limitations, including poor access to ancient genomes and genome-wide markers (44–48).

Here, we publish the first whole-genome data on Atlantic bluefin tuna including a long time-series of genome-wide aDNA data, analysing 41 archaeological and archived bones from seven Mediterranean and Norwegian locations, dated from ca. 3000 BCE to 1941 CE (Figure 1). Using whole-genome sequencing data of a further 49 modern individuals caught in the Mediterranean, Norway and the Gulf of Mexico between 2013-2018, we investigate population admixture, effective population size and genetic diversity over time to test for the impact of overexploitation on bluefin tuna demography.

## Results

We analysed over 19.8 billion sequencing reads, obtaining an average of 9.9-fold nuclear coverage for the ancient and 11.9-fold coverage for the modern samples (Table 1, S1). Ancient samples showed excellent preservation with relatively high proportions of endogenous DNA (average 40.4%), while containing fragmentation and sequence damage patterns (increases in C>T and G>A transitions) that are expected from authentic, degraded DNA (Table 1, Fig. S1). We mapped all reads to a pseudo-chromosome reference genome produced from high-quality, short-read data for the species and—following an extensive set of quality-filtering steps (Materials and Methods)—we analysed single-nucleotide polymorphisms (SNPs) in 90 individuals using approaches based on hard called genotypes. Overall, these filters produced final pruned datasets of 386,963 loci for modern spawning area samples and 12,838 loci when combining modern and ancient samples. This represented a ca. 15-fold increase in genomic representation compared to previous attempts investigating modern BFT demography (33), and over a 100-fold increase when using ancient DNA (45, 49).

**Table 1.**
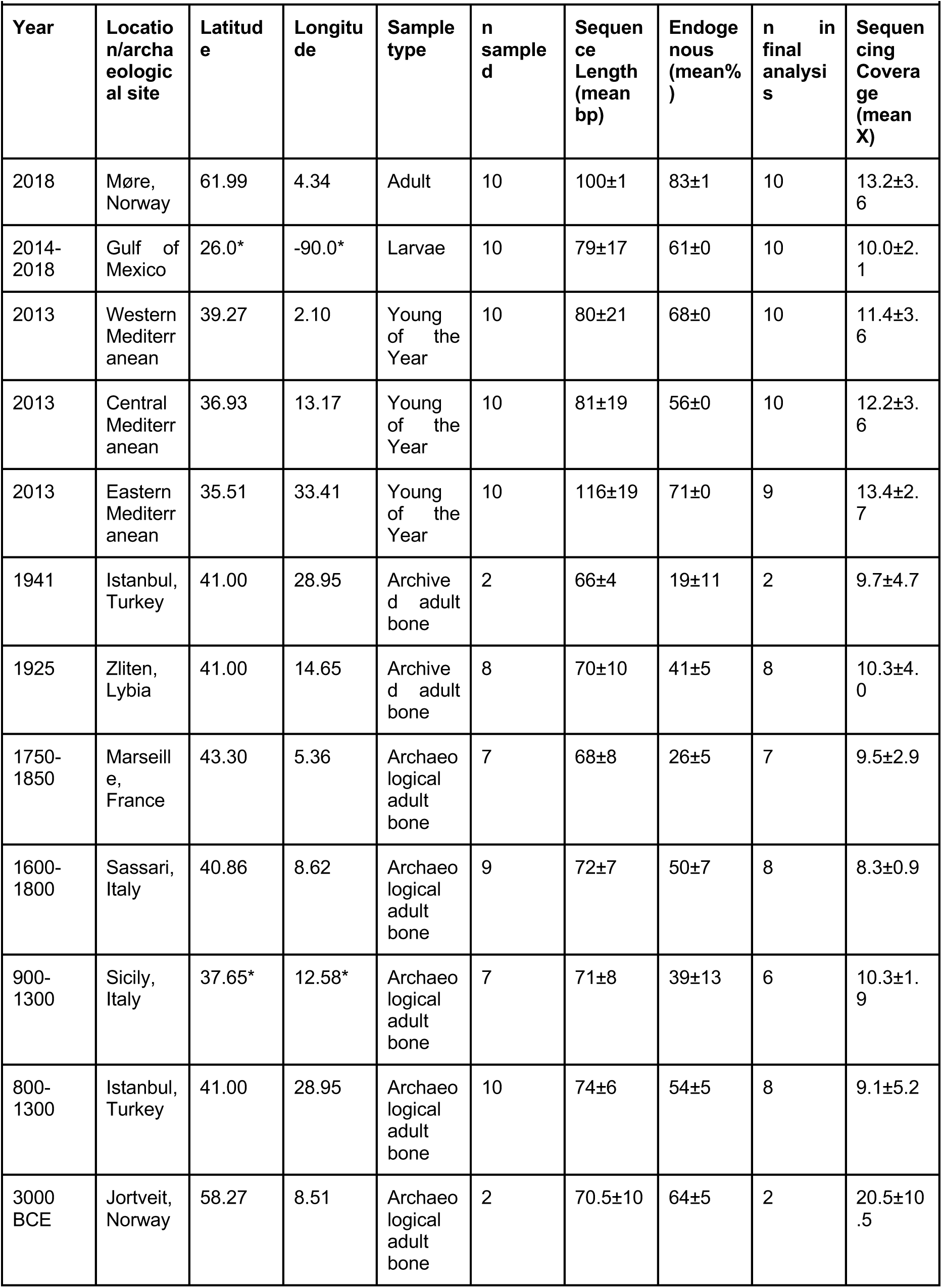
Summary details of ancient and modern Atlantic bluefin tuna samples sequenced herein. n = number. Gulf of Mexico and Sicily samples pertain to several sites, for full details see Supplementary Table S1.

### Patterns of population mixing consistent across millennia

We found that the most likely number of bluefin tuna populations (K) described by PCA and STRUCTURE was two, represented by one genomic cluster for the Gulf of Mexico, and one for Norway and the Mediterranean (Fig. 1B-D, Fig S2). We found increased (ca. 10%) western-Atlantic ancestry in individuals caught off Norway and in the Eastern Mediterranean - which was consistent in ancient samples dating back up to 3000 BCE from the same areas (Fig. 1C), and at 386,963 loci for the modern spawning site dataset (Fig. S3). Bluefin tuna from Western and Central Mediterranean sites, over a millennium, shared similar ancestry proportions to each other, with lower Gulf of Mexico-like ancestry and increased Mediterranean ancestry than Eastern Mediterranean individuals (Fig. 1C).

Pairwise F_ST_ values (Fig. 1D) across 386,963 loci for modern spawning sites support admixture patterns where Eastern Mediterranean individuals are slightly more closely related to Gulf of Mexico individuals than the Western and Central Mediterranean are (0.0017 vs 0.0018 F_ST_). Overall, differences between the Western and Eastern Atlantic are low but significant (p=0.007). Patterns of population structure appear unbiased by our loci filtering choices on both modern and ancient datasets (Fig. S3, S4). Our results were not driven by recently-discovered introgression, which was identified on Chromosome 8 and was filtered out (along with other deviations from HWE, See Methods, Fig. S5, S6) on the basis that they are not population-defining characteristics (33) and present only in a few individuals (data not shown).

### A historical demographic decline and genetic erosion

Reconstructions of historical effective population size (N_e_) using GONE reveal significant N_e_ decreases in both the Gulf of Mexico and Mediterranean spawning area samples between 1850-1900 (Mediterranean: p<0.001, R2=0.15, F=27.7_149_ linear regression), corresponding to a period of intensive exploitation. Runs of homozygosity (RoH) show inbreeding extent (fraction of genome in RoH, FRoH) varied slightly between modern spawning areas where inbreeding appears greatest for Western and Central Mediterranean samples (Fig. 2B), and the Eastern Mediterranean sample is slightly significantly lower (p<0.05 t-test), with the Gulf of Mexico an intermediate group.

**Figure 2.**
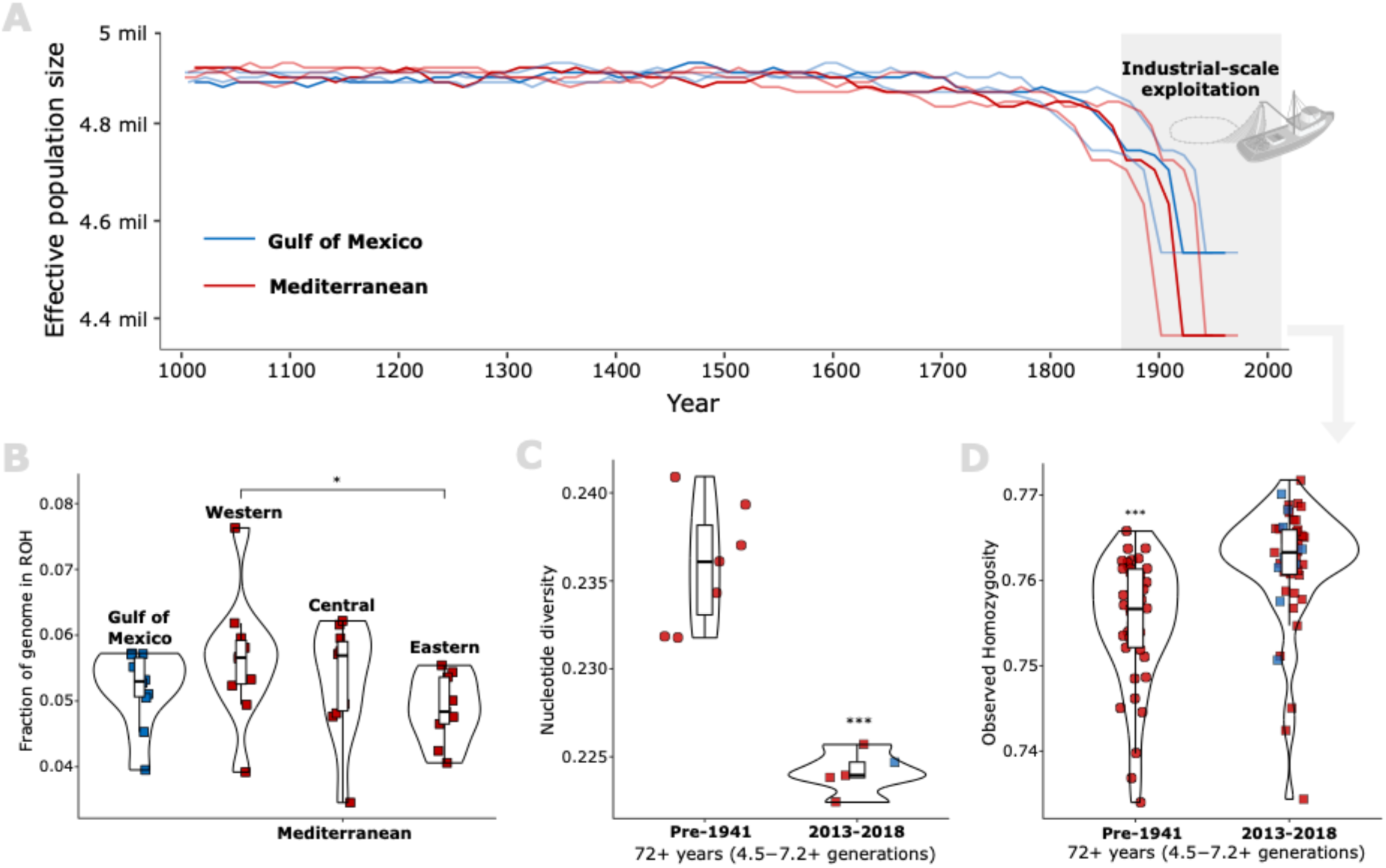
Temporal demographic decline and genetic diversity loss in modern Atlantic bluefin tuna (*Thunnus thynnus*). (A) GONE reconstructions of historical effective population size (Ne) for the Gulf of Mexico (blue, n=10) and Mediterranean (red, n = 29) populations using modern spawning site samples at 664,522 SNPs. Years for GONE estimates were calculated from generation numbers using three different generation lengths (10-16 years), showing the most likely (13 years) in full colour, and 10 and 16 years slightly transparent. GONE plots do not include data more recent than 1973 due to the removal of estimates for the four most recent generations (see Methods). Estimates were illustrated in relation to a potential causative exploitation event. (B) Violin plots show the fraction of the genome in runs of homozygosity (RoH) for each modern spawning area sample, at 664,522 unpruned SNPs. (C.) Violin plots show nucleotide diversity (per sample group) differences at 2,890,241 SNPs including 10% missing data between pooled ancient (Pre-1941, sample groups=7) and modern (2013-2018, sample groups=5). (D) Violin plots show observed homozygosity (fraction of sites) differences at 17,741 SNPs between pooled ancient (Pre-1941, n=41) and modern (2013-2018, n=49). Significance of t-tests between the groups was illustrated for significant pairs using asterisk (***: p<0.0001, *: P<0.05), boxplots illustrated with group means, 25^th^ and 75^th^ percentile as outer edges and 95^th^ percentiles (black whiskers). Samples are coloured by population (blue: western stock, red: eastern stock) where shapes depict time period (circle: ancient, square: modern).

Modern samples (2013–2018) had significantly lower nucleotide diversity and higher homozygosity (lower heterozygosity) than ancient samples (3000 BCE - 1941 CE, p<0.0001 t-test, Fig. 2D,E). Nucleotide diversity and homozygosity results show these are genome-wide patterns i.e. tens of thousands of loci spread across the genome but are subject to missing data at 2,890,241 loci (Fig. S7, S8). The loss of heterozygosity appears to have begun from the period of 1600-1800 (years) and declined step-wise, again after 1941 (Fig. S9). Low sample size per group and spatial variance across the Mediterranean complicate inferences of a temporal trajectory but the same general stepwise trajectory of loss in nucleotide diversity is observed (Fig. S10).

Population-specific F_ST_ was analysed as a measure of expected heterozygosity which is theoretically less impacted by mutation and inbreeding than RoH (50, 51), where the most heterozygous sample is theoretically most ancestral (52). Population-specific F_ST_s are lowest (most heterozygous i.e. ancestral) for the Gulf of Mexico, which shares overlapping confidence intervals (CIs) with the Eastern Mediterranean, while the Central and Western Mediterranean (sharing overlapping CIs) have higher F_ST_s (Fig. S11), corroborating STRUCTURE and FRoH results. The Gulf of Mexico sample overlaps more with the Mediterranean samples in inbreeding extent (FRoH) than population-specific F_ST_ (Fig. 2C,D). Lengths of RoHs were relatively short (<175 Kb) and did not vary between modern spawning sites (Fig. S12, S13).

The Gulf of Mexico sample and Mediterranean samples share similar N_e_ over millennia, beginning (from year 1000) with a period of long stability across centuries, before a significant decline around year 1850-1900, regardless of whether the Mediterranean samples are pooled (Fig. 2A, Fig. S14). Significantly higher homozygosity for modern samples was consistent when removing transition sites which had the potential to be affected by post-mortem DNA damage (Fig. S15, p<0.001 t-test). Modern samples also had higher overall homozygosity than ancient samples if we did not filter loci by F_ST_ and if we allow missing data (Fig. S8, p<0.001, t-test). F_ST_ differences between ancient and modern samples show that temporally variable loci are relatively low F_ST_ (<0.1) and do not display patterns across the genome indicative of selective sweeps (Fig. S7).

## Discussion

Here, we used ancient and modern whole-genome data of Atlantic bluefin tuna to assess genomic consequences of their exploitation, such as changes in population structure, admixture and genetic diversity. We find evidence for two populations: a western and eastern Atlantic component that share high levels of connectivity, especially in bluefin tuna caught off Norway and the Eastern Mediterranean. We find genomic evidence that bluefin tuna population(s) suffered a demographic decline by 1850-1900, and we correlate this observation with a loss in heterozygosity over the same period. Here, we contextualise our results with ecological, evolutionary and historical perspectives.

### Patterns of population mixing consistent across millennia

Our study suggests bluefin tuna population structure has persisted across millennia, comprising two populations with low levels of differentiation. This finding is consistent with the ecology of bluefin tuna as a highly fecund batch spawner exhibiting wide-ranging migrations, high levels of population mixing, and large census sizes, as well as previously-published genetic results (24, 25, 53), (26, 33). Low genetic divergence in bluefin tuna complicates the assessment of population structure. Nevertheless, we find small (ca. 10%), increased western Atlantic-like ancestry in Norwegian and Eastern Mediterranean samples that are consistent across millennia suggesting that these are true biological patterns.

These admixture patterns may be ecologically explained if bluefin tuna caught off Norway and in the Eastern Mediterranean mix with Western Atlantic bluefin tuna. A phenomenon we find likely in poorly understood Atlantic spawning areas, like the Slope Sea and Bay of Biscay or potential ones like the Azores and Canary Islands (Fig. 1; (25, 28, 31)). It is known that a portion of the Mediterranean bluefin tuna spawn with Western Atlantic bluefin tuna in the Slope Sea (33). We suggest this phenomenon may be widespread in other under-studied spawning areas and vary between Mediterranean bluefin tuna. While we cannot exclude the possibility that our results are due to error or small sample size, we find it unlikely that either can explain the temporal continuity in admixture through samples of various origins. We suggest that variation in bluefin tuna admixture within the Mediterranean was probably not found in previous studies due the pooling of Mediterranean samples (33), or using small SNP datasets rather than whole genome sequencing done herein (26, 27). Furthermore, greater genetic connectivity between the Atlantic and Eastern–rather than Western–Mediterranean spawning grounds is a component of other marine fishes (54–56), potentially driven by a spawning behaviour established when Atlantic species re-colonised the Mediterranean at the end of the last glacial maximum (57).

Theories of resident and migratory components of Mediterranean bluefin tuna have existed for almost as long as the period our dataset covers (6, 34, 35). Our findings do not suggest that Western and Central Mediterranean bluefin tuna are resident, rather that bluefin tuna that spawn at these locations are less likely to have originated from Atlantic spawning sites or the eastern Mediterranean. Tagging studies document well that bluefin tuna caught in the Western and Central Mediterranean during the spawning season migrate to the Atlantic (34, 58, 59). Studies tagging Norwegian or Eastern Mediterranean spawners are rare (35) and thus it is possible that bluefin tuna from these locations may be more resident in the Mediterranean in some years, as isotope data across millennia suggests (37), while in other years contributing more to Atlantic spawning than Western and Central Mediterranean bluefin tuna. We find the latter most likely since greater variation in spawning behaviours and increased mixing rates with the Western population are mechanisms, which maintain heterozygosity as observed in Eastern Mediterranean bluefin tuna.

There is a lack of knowledge on the contribution of each stock to spawning at Atlantic spawning sites, other than for the Slope Sea – where a mix of the two stocks was evident (33). We theorise increased spawning rates of Eastern Mediterranean and Norwegian bluefin tuna than Western and Central Mediterranean bluefin tuna in Atlantic spawning areas. A host of research supports increased complexity in bluefin tuna migrations and spawning than is currently recognised (6, 28, 33). For example, a proportion of bluefin tuna are genetically Mediterranean–but isotopically Western Atlantic (60), and a proportion of bluefin tuna may engage in skipped-spawning (61). Further work is required to assess whether Norwegian and Eastern Mediterranean bluefin tuna spawn in the Atlantic, while being more resident for the years that they spawn in the Mediterranean, including in the Black Sea as has been hypothesised (62, 63).

Depending on the season, the geographical and genetic distance between Mediterranean spawning sites could be minimal (25). Therefore, the high likelihood that spawning segregation is not always present in the Mediterranean could explain why the genetic differences between Western and Central vs. Eastern Mediterranean are not significant, despite that the admixture appears consistent in snap-shots across time. Due to the limited extent of these admixture patterns, caution should be taken when interpreting our findings and further research should be done to investigate the full extent and drivers of differences in spawning site use in Mediterranean bluefin tuna, whether they may be behavioural (migrational memory) (64) or genetic.

### A historical demographic decline and genetic erosion

We find genomic evidence that bluefin tuna began a significant demographic decline by the beginning of the 1900s, which is earlier than robust fisheries data became available (ca. 1960, (4), and therefore was previously unknown. Currently, quantitative data recognises a demographic decline between 1960-2007, where Eastern bluefin tuna abundance and range declined by an estimated 70% and 46-53%, respectively (4, 5), followed by a recent recovery (13). Our demographic modelling suggests that the post-1960 losses represent only the most recent impacts of intensive exploitation. Supporting this, a host of evidence would point to exploitation being more intensive and impactful earlier than currently recognised. These include declines in prey and habitat suitability by the 1800s (16, 37), the highest tuna trap catch rates in history by the 1880s (40), a shift to fishing larger individuals by the early 1900s (39), and the seasonal, geographic and gear expansion of fishing into the Atlantic by the early 1920s (3, 15). Together with findings on other species like Atlantic cod (*Gadus morhua*) (65) and herring (*Clupea harengus)* (*16*), pre-industrial demographic declines appear to be common for commercial Atlantic marine fishes.

Since bluefin tuna fisheries seldom existed in the Western Atlantic until the 1950s (3), it is unlikely that Gulf of Mexico bluefin tuna declined to the same extent as the Mediterranean, as our data suggest. Similar demographic histories for both populations over the past millennia could be due to the limitation of the measure of effective population size, which is affected by gene flow between populations (33, 66). Effective population size thus cannot be interpreted as biomass for each population but rather the increased variance in reproductive success as a probable result of a biomass decline of the species (67). Given that bluefin tuna demography appeared stable across hundreds of years prior to the 1800s, we suggest that this increased variance in reproductive success was caused by novel challenges like increased fishing effort, causing more dramatic fluctuations in biomass than is normally reflected by climate and other factors (6, 38, 68).

A widely accepted consequence of biomass declines is genetic bottlenecks, in which genetic diversity decreases due to a loss of alleles and increase in inbreeding. This has been well documented for other taxa such as marine seabirds (43), but evidence has been scarce for fish populations. Our work suggests that marine fishes are not immune to genetic erosion following the finding of decreases in heterozygosity and nucleotide diversity, corresponding to a period of overexploitation and demographic decline by the early 1900s. The only other whole genome aDNA analysis on marine fishes showed genetic diversity retention in the overexploited Atlantic cod (44), though this study did not analyse samples older than the year 1900. Both genetic stability (45, 46, 49, 69–71) and erosion (47, 72, 73) have been observed using fewer SNPs, microsatellites or mitochondrial DNA. Our findings do not suggest that bluefin tuna is at risk of extinction. The genomic changes we observe are relatively small and a host of information would lead to evidence that its populations are rebounding (9–11, 13).

It is likely that a combination of bluefin tuna ecological features buffer against losses in genetic diversity such as high population connectivity, large population size and a long life cycle which promotes heavily overlapping generations (25, 74). Theoretically, each of these features limit the effect of genetic drift (75, 76), such that a larger number of generations would need to have elapsed in order to observe changes in heterozygosity (R. Waples, pers. comm.) as a result of biomass declines. This may explain why the decrease in heterozygosity and nucleotide diversity we observed between ancient and modern bluefin tuna was slight (ca. 1-1.5% sites), though significant. Dictating the degree of genetic diversity loss in modern samples, we find that inbreeding extents (fraction of the genome in runs of homozygosity) differ between modern spawning areas, which corroborate patterns of admixture i.e. Eastern Mediterranean bluefin tuna had a significantly lower inbreeding extent than Western Mediterranean bluefin tuna, which we hypothesise is due to a greater contribution of Western Atlantic gene flow to the Eastern Mediterranean.

There is concern that the Western Atlantic population is at genetic risk due to a perceived greater population decline coupled with a strong recovery in the Mediterranean population (13, 33). Our work suggests that the Western Atlantic and Mediterranean populations do not significantly differ in inbreeding extent or nucleotide diversity. Overall, population-specific F_ST_ indicated that the Gulf of Mexico is more heterozygous than the bluefin tuna of the Mediterranean spawning sites as has been suggested previously (33), potentially being more ancestral–and thus less impacted. We agree with (33) that an increasing number of east to west migrants (77) may have the potential to erode differences between the Western and Eastern stocks, but our results suggest that the mixing of the populations in Atlantic spawning areas has been a phenomenon for millennia.

### Approach and limitations

Due to ancient sample availability and post-mortem DNA damage, we here opted to study a high-quality yet relatively small dataset in terms of sample size and SNPs. This is a vast improvement on studies that focused on only a handful of SNPs, but could still be improved in further studies that allow for deeper sequencing and an improved genomic reference. In addition, we filtered SNPs by F_ST_ to remove highly-conserved or erroneous sites when analysing ancient samples, as these sites – likely to be under balancing selection – have been shown to reduce the appearance of population differentiation (76,77) (Fig. S4). Low F_ST_ sites are likely to maintain genetic diversity in marine populations (78); when including these loci we found that proportions of heterozygosity loss were lower over time (Fig. S8). We consider that the inclusion of highly-conserved loci would have hindered our observations and is not always appropriate for detecting changes in datasets with low levels of population divergence and ancient DNA damage. On the other hand, certain analyses e.g. nucleotide diversity required sufficient window SNP density such that a conservative approach was not possible with our dataset where we had to allow missing data into analyses. The variety of filtering methods taken provide support that patterns observed are likely to be true and appear regardless of approach.

Any study investigating effective population size should reiterate that its relationship with census size is complex for fishes such as bluefin tuna with very large census sizes (76). Moreover, effective population size estimates are influenced by many factors (67, 79), chiefly gene flow–which in the case of bluefin tuna appears high. Due to a lack of pre-industrial Western Atlantic overexploitation, we propose that gene flow from the Western stock buffered the demographic decline we observed in bluefin tuna by 1900. The timing of the decline is biased by the chosen generation length. In this study, we used three alternatives to be conservative, but their consistency and relevance over time is unknown. Limitations of the method also precluded the opportunity to realise effective population sizes for the past ca. 40 years (ca. 4 generations), which should be explored to capture the full extent of the demographic decline and contextualise the recent recovery.

Our lack of a high-quality annotated reference precluded the opportunity to assess the location of SNPs which vary in space or time and their function. Future studies should assess gene ontology and assess mechanisms underlying these polymorphisms. For example, they may represent polygenic or epistatic, global adaptation (75, 80–83), even if they are at low-F_ST_ differences in space or time.

Aided by ancient DNA, we provided directions for future research by indicating that the spawning, recovery and genetics of bluefin tuna is likely to be more complex than currently recognised. Improved genomic resources are almost certain to facilitate improved understandings on the scale of impacts and diversity of populations, especially in cases such as bluefin tuna where population differentiation is low. Our findings call for additional analyses on temporal analyses of bluefin tuna and other marine fishes, using high-quality annotated reference genomes, larger sample sizes and coverage. We show how ancient DNA has the potential to aid marine management and conservation plans by providing novel perspectives of when demographic declines began to revise recovery targets and novel ways to measure population sustainability, depending on what genetic diversity is lost and its functional implications.

## Methods and Materials

### Sample collection

We collected samples of modern bluefin tuna for analysis as follows: contemporary larvae or young-of-the-year specimens (Gulf of Mexico, Western Mediterranean - Balearic Islands, Central Mediterranean - Sicily, Eastern Mediterranean - Levantine Sea, n = 40, Table 1; Table S1) were collected from each of the major BFT spawning sites between 2013 and 2018 (Fig. 1, Table 1; Table S1). In addition we collected and analysed adult bluefin tuna (n=10) caught off the Norwegian coast in 2018. For further sample details see Supplementary. Larvae or muscle tissue samples from each specimen were preserved in 96% ethanol and stored at -20 °C until extraction.

Ancient samples included archived vertebrae (n = 12; Table 1) which were retrieved from the Massimo Sella Archive (49) and pertained to two tuna-trap catches of 1941 (Istanbul, Turkey) and 1925 (Zliten, Libya: Table 1, Fig. 1A). Ancient samples also included archaeological vertebrae (n = 32, Table 1) retrieved from several Norwegian and Mediterranean excavations throughout the past five millennia (Fig. 1A) including 1750-1850 (Marseille Harbour, Marseille, France), 1600-1800 (Pedras de Fogu, Sassari, Italy), 900-1200 (Sicily, Italy), 800-1300 (Yenikapi, Istanbul, Turkey), and 3000 BCE (Jortveit, Norway). See Supplementary Materials 1 for more details on ancient samples and their dating.

### Ancient DNA extraction

The majority of archaeological and historical bluefin tuna samples underwent ancient DNA (aDNA) extraction in dedicated, sterile, PCR-free conditions at the Ancient DNA Laboratory of the Department of Cultural Heritage (University of Bologna, Ravenna Campus, Italy). The two ancient Norwegian samples were processed in a dedicated aDNA laboratory at the University of Oslo as detailed in (84). Samples processed in the Ancient DNA Lab followed strict criteria for aDNA analysis (85, 86). The outer surfaces of bones were cleaned with a ca. 20% sodium hypochlorite (bleach) solution and left to air-dry for 10 minutes. Specimens were then exposed to UV light (254 nm) for 15 minutes on each side before drilling to remove an outer layer (ca. 2 mm) of material. Between 100-350 mg of bone powder was then collected by drilling at the same position where the outer layer had been removed. Care was taken to avoid overheating specimens by drilling at slow speeds with diamond-tipped drill-bits.

Isolation of aDNA was performed using a modified version of Dabney et al. (87, 88). Briefly, 100-300 mg bone powder from each sample was pre-digested (89) for 20 minutes at 37°C in 1-3 mL digestion buffer containing EDTA (0.45 M, pH 8.0) and proteinase K (0.25 mg/mL). Lysates were then discarded before samples were fully digested overnight in the same conditions. Once digested, lysates were combined with 10 mL of binding buffer (PB buffer, Qiagen, Germany) and bound to the High Pure Viral Nucleic Acid Large Volume silica spin columns (Roche, Basel, Switzerland) by centrifugation. Membrane-bound aDNA was then washed twice with 720 µL PE buffer (Qiagen), before elution in 30 µL of EB buffer (Qiagen). The total DNA obtained from each extraction was quantified using the Qubit dsDNA HS (High Sensitivity) Assay Kit (Thermo Fisher Scientific, USA). Negative controls employed in each batch of samples extracted indicated an undetectable level of contamination (<500 pg/mL). Extractions were then stored at -20°C until library preparation.

### Modern DNA extraction

Modern spawning site specimens were extracted at the GenoDREAM laboratory of the Department of Biological, Geological and Environmental Sciences (University of Bologna, Ravenna Campus, Italy). Modern Norwegian samples were collected by the Norwegian Institute of Marine Research (IMR), freeze dried and powdered at IMR facilities and extracted in the modern DNA isolation laboratories at the Center for Ecological and Evolutionary Synthesis, University of Oslo, Norway) as detailed in (84). Modern spawning site samples were processed using a modified salt-based extraction protocol (90) using SSTNE extraction buffer (91), and treated with RNase (Qiagen) to remove residual RNA. After isolation, the total DNA obtained from each extraction was quantified using the Qubit dsDNA BR Assay Kit (Thermo Fisher Scientific, USA). Negative controls employed for each batch of samples extracted confirmed an undetectable level of contamination. Samples were diluted to 10 ng/µL. Prior to library preparation, 100 µL of DNA extracted from each modern sample was sheared to maximum sizes of 500 bp using a Bioruptor Pico sonicator (Diagenode) with the settings: Medium 30 on/90 off for 10 minutes. Samples were precipitated using isopropanol, following a procedure from Qiagen (FAQ-ID-2953). Fragment lengths were confirmed by agarose gel electrophoresis and total DNA was re-quantified using Qubit before precipitates were stored at -20°C until library preparation.

### DNA library preparation and sequencing

With exception to the Norwegian samples which were prepared as detailed in (84), single-stranded libraries were built for both ancient and modern samples using the Santa Cruz Reaction (SCR) method (92) from 10-20 µL of DNA, up to a maximum input of 150 ng. Prior to indexing, aDNA libraries were prepared in sterile conditions at the Ancient DNA Laboratory of the Department of Cultural Heritage, separate from those facilities used to prepare modern libraries. Library quality and the non-amplification of controls were confirmed using qPCR following the SCR method, which indicated the number of cycles required for indexing and inhibited samples to be discarded. Ligated DNA was double-indexed with sample-specific 6 bp indexes by amplification using 2X Amplitaq Gold 360 MM (Thermo Fisher Scientific, USA), and 25-50 µM of each index for 10-14 cycles (2 min at 95°C, cycles of 30 s at 95°C, 30 s at 60°C, and 70 s at 72°C with a final extension of 10 min at 72°C). Amplified products were cleaned by using AMPure XP beads (Agencourt) at a 1:1.2 ratio, eluted in 25 µL in EB buffer and quantified using a Bioanalyzer 2100 (Agilent Technologies). All libraries were screened to explore DNA preservation and library clonality by sequencing ca. 1 million reads per sample on the Illumina HiSeq X, 2500 or 4000 platforms (100 bp paired-end). Libraries were then re-sequenced (150 bp paired-end) on the HiSeq 4000 or NovaSeq S4 Illumina platform, to the final sequencing depths reported. Sequencing and demultiplexing (allowing zero mismatches in the index tags) were performed at the Norwegian Sequencing Center (Oslo, Norway) and Macrogen facilities (Seoul, South Korea/Amsterdam, Netherlands) with care taken to avoid batch effects by pooling varied combinations of samples (Table S1).

### Reference assembly improvement

Since a chromosome-level assembly was not available at the time of analyses, we improved a draft assembly. The highly-scaffolded draft *Thunnus thynnus* genome assembly (NCBI BioProject: PRJNA408269) was mapped against the chromosome-scale assembly of yellowfin tuna (*Thunnus albacares*, fThuAlb1.1, NCBI accession: GCA_914725855.1). Scaffolds were mapped using BBMap (93) with the asm10 setting which successfully mapped 102,121 out of 103,645 (98.5%) scaffolds to yellowfin tuna chromosomes. Scaffolds were then binned into the 24 yellowfin tuna chromosomes by placing 200 N’s between them, resulting in a 768 Mb pseudo-chromosome assembly. Non-mapped (unplaced) scaffolds were excluded from analyses.

### Data processing and filtering

Due to initial read lengths being longer in modern Norwegian samples (ca. 160 bp, Table S1), because DNA was not sonicated as modern Gulf of Mexico and Mediterranean samples, we first trimmed Norwegian reads to 100 bp. Our aim was to produce a dataset that was technically comparable and avoid one sample group having sequence length or depth much greater than others. Raw Norwegian sequences were trimmed using Trimmomatic v.0.39 (94), using the settings clip=100, headclip=5. Details of average sequence lengths across all samples can be found in Table S1.

Ancient and modern reads were processed by using PALEOMIX (95) which is a set of pipelines and tools designed to aid the rapid processing of high-throughput ancient sequencing data. Forward and reverse reads were collapsed in PALEOMIX with AdapterRemoval v1.5 (96) and aligned to our pseudo-chromosome BFT reference using BWA mem (93). Only reads that had a minimum length of 25 bp were aligned, and only with a minimum quality score (MapQ) of 30 i.e. a 0.1% chance each read was mis-aligned were used for subsequent analyses. Ancient DNA damage patterns were investigated by using mapDamage v.2.0.6 (97). Finally, we removed all clipped reads (or pairs of reads for which one read was clipped) and all duplicate reads (Picard Tools v. 1.96) and then realigned indels using GATK v. 3.7 IndelRealigner (98). To reduce the influence of post-mortem taphonomic degradation on our downstream SNP analyses, we studied damage patterns (Supplementary Figure S1) and trimmed 3 bp from all ancient mapped reads using the TrimBam function of bamUtil v.1.0.6 (99) and re-indexed using samtools v1.7 (100). This reduced the frequency of deamination to below 5% in all reads. Average read-depths were 9.9 and 11.9 fold coverage for ancient and modern samples, respectively.

We used GATK to jointly call SNPs (GATK HaplotypeCaller and GenotypeGVCFs) for ancient and modern samples using default settings, allowing a maximum of three alternative alleles. We filtered SNPs using conservative established approaches. First, SNPs were hard-filtered for strand bias (FS and SOR), mapping quality (MQ), quality by depth (QD), non-polymorphism (AC) using BCFTOOLS v. 1.6 with settings filter --i ’FS>60.0 || SOR>4 || MQ<30 || QD<2.0 || AC==0 || AC==AN’. Then, we removed indels and retained only, bi-allelic SNPs using VCFtools v.0.1.16 (101) with settings --remove-indels --min-alleles 2 --max-alleles 2.

VCFtools was used to filter for genotype quality (GQ), depth (minDP and max-meanDP) and minor allele frequency (MAF). A minimum GQ of 20 was applied to remove low confidence genotypes expected due to aDNA characteristics such as post-mortem damage and highly variable coverage across the genome. SNPs present in genomic regions where reads preferentially mapped more than twice the average (likely repetitive regions) were removed using a max-meanDP filter set to 30; ca. double the coverage of the majority of our highest coverage samples. A minDP filter of 5 was used, to provide confidence that homozygous loci were not artefacts of insufficient coverage. MAF filters were used to remove doubletons which may represent poor sequencing or DNA quality, and set depending on the number of samples present in multi-sample VCFs i.e --maf 0.05 or 0.02 if using modern (n=50) or modern and ancient (n=95) samples, respectively. No missing data was tolerated for analyses (--missing-data 1). Finally, to retain sufficient loci with this conservative approach, three ancient samples were dropped from analyses due to low coverage <6X (IST_913C_01, IST_913C_02 and PF_1618C_23, Table S1).

We excluded loci that were out of Hardy-Weinberg equilibrium (HWE) due to the possibility that highly hetero-or homozygous loci represent non-biological sequencing artefacts, or signatures of introgression that could be present in few individuals (e.g., (33) and may bias analyses with small numbers of samples. We opted to remove loci if they yielded a p-value <0.001 following a site-specific exact test across all samples as implemented in VCFtools. The trade-off with this filter is that we will not observe highly variable sites which may be signatures of selection however since recently no large effect loci have been observed in bluefin tuna (33), we assumed that loci that were highly hetero- or homozygous are more likely non-biological sequencing artefacts. We illustrate the effect of HWE filters in the supplementary (Figure S5, S6).

Due to the possibility that multiple ancient bones were sequenced from the same individual, or that modern tissues were sampled from closely-related individuals that might bias analyses, we tested pairwise relatedness between samples in VCFtools using -- relatedness2. This resulted in the removal of two ancient specimens from analyses (CDM_10C_07 and CDM_10C_11) due to their kinship coefficient >0.35 i.e. duplicate/twin with CDM_10_04. The modern sample CYPR-LS-331 was removed for the same reason due to its close-kinship >0.35 with CYPR-LS-330.

Overall, these filters produced final datasets across the 768 Mb bluefin reference genome of 39 modern spawning site individuals with 664,522 loci and combined ancient and modern datasets of 90 individuals with 30,264 loci.

To remove highly conserved or erroneous sites from the dataset which did not vary consistently over time or space and that therefore may have clouded inference of spatio-temporal variability, we assessed F_ST_ per site and filtered out loci if their F_ST_ values were <0 across all spatio-temporal sample groups. Setting the filtering threshold at <0 F_ST_ was an arbitrary but conservative approach, removing sites that were on average more variable within groups than between them. This approach was preferred to filtering and analysing ‘outlier loci’ as is common practice for fishes with low levels of population differentiation e.g. (102, 103) since population differentiation or adaptation in bluefin tuna is most likely not driven by large effect loci (33) and therefore these loci were perceived as errors, likely not-population defining. Therefore our aim was to retain relatively low F_ST_ but consistently (between time periods) variable sites in the dataset. F_ST_ was estimated in VCFtools with the commonly used --weir-fst-pop function (104, 105) with default settings. This function calculated per-locus pairwise F_ST_ (50), between all spatio-temporal groups. A distribution of F_ST_ per locus from the calculations can be observed in the supplementary (Fig S16). VCFtools was also used as the means to filter out loci using the --position function. F_ST_ filtering resulted in 17,741 loci of the 30,264 loci remaining.

### Population structure and admixture

SNPs were pruned (–indep-pairwise 50 5 0.2) for linkage disequilibrium using PLINK v1.90b6.21 (106) following (107). Therefore, analyses were performed on two datasets; one pruned modern dataset of 386,963 loci, and one filtered, pruned modern and ancient dataset of 12,838 loci. A principal component analysis (PCA) was performed using EIGENSOFT v.5.1 (108). PCA is a method free of HWE assumptions but is open to technical bias (109), therefore we opted to avoid overinterpreting these results. We re-ran analyses and included sites previously removed by our F_ST_ filter threshold (F_ST_ >0) and re-ran PCA analyses on 27,882 loci to explore the need for filtering loci by F_ST_ to observe and infer correct population structure.

Population structure was also evaluated on the same datasets using STRUCTURE v.2.3.4 (110), which implements a Bayesian clustering method assuming HWE to identify the most likely number of populations (K). We followed the Evano *et al*. (111) method, and thus, we carried out 10 runs per each value of K ranging from 1 to 5. Runs used the admixture models and assumed correlated allele frequencies without locprior. Each run used 10,000 burn-in and 50,000 Markov Chain Monte Carlo replicates. We estimated the ad hoc statistic ΔK in order to infer the most likely number of populations using STRUCTURE HARVESTER (112). CLUMPAK (113) was used to merge the 10 runs from the most probable K, which reported similarity scores >95%. Pairwise distances between modern spawning site samples were calculated with Nei’s estimator of F_ST_ (114) as implemented in the hierfstat R package, using 1000 permutations to calculate p-values, which were judged for significance under the FDR approach at the 5% level.

### Genetic diversity statistics

Genetic diversity statistics were calculated using the filtered, dataset of 90 modern and ancient samples at 17,741 loci. Heterozygosity (proportion of sites) was calculated for each individual using VCFtools --het. We removed transition sites (C > T and G > A SNPs) which may be affected by post-mortem taphonomic degradation and re-ran the heterozygosity analysis on the remaining 5822 loci, which revealed our findings hold true irrespective of the potential DNA damage. Finally, we removed the per-site F_ST_ filter and allowed 10% missing data (using VCFtools --missing-data 0.9) into the dataset to produce 2,890,241 loci and re-ran analyses which revealed that these patterns appear to be genome-wide. The same dataset, reprocessed using BCFTOOLS to include non-variant sites, was used to calculate nucleotide diversity (pi). Nucleotide diversity (pi) was calculated for each sample group per 50kb (-w 50000 -m 10) windows using the script popgenWindows.py (https://github.com/simonhmartin/genomics_general). As a proxy for inbreeding extent, runs of homozygosity (RoH) were analysed for the modern spawning site samples using a filtered unpruned dataset of 664,522 loci following (115). ROHs were calculated using PLINK across 100kb windows following (16) with the following settings homozyg-snp 20 --homozyg-kb 100 --homozyg-density 20 --homozyg-gap 1000 --homozyg-window-snp 20 --homozyg-window-het 2 --homozyg-window-missing 5 --homozyg-window-threshold 0.05. T-tests were performed in R to judge the statistical significance of summary statistic differentiation between sample groups, at the 5% level.

### Effective population size

We reconstructed historical effective population size (N_e_) using GONE v.1 (116). GONE uses linkage disequilibrium (LD) to estimate N_e_ for the previous 200 generations. GONE was run on the unpruned filtered dataset of modern spawning site samples comprising 664,522 loci, grouped by western or eastern stock (Gulf of Mexico, n = 10; Mediterranean n = 39). We converted GONE outputs of N_e_ estimates per generation number to years (BCE/CE) using a generation length of between 10 and 16 years, with 13 years being the most widely accepted following (76) and https://fishbase.se. We discarded estimates of the previous 4 generations (ca. last 50 years) as these yielded extremely low N_e_ estimates (ca. 15-20k), likely as a function of methodological limitations (79). We re-ran analyses after removing the maf filter which resulted in similar trajectories and tested for the effect of sample-size by grouping Mediterranean samples by spawning site (n = 9 or 10), which yielded near-identical N_e_ trajectories over time as when analysed pooled (Fig. S11).

### Outlier analyses

To assess the impact of HWE filters and investigate outliers across the genome which we hypothesised would include patterns of introgression recently observed in bluefin tuna (33), we used the SNP-based outlier detection method with the lowest false positive rates which detects outlier loci using allele frequencies; pcadapt (117). pcadapt does not require sample groups to be defined. We ascertained population structure via PCA and determined outlier loci as those excessively correlated with two (K=2) ordination axes. pcadapt was run in R using the unpruned HWE-filtered and non-HWE-filtered modern spawning site datasets of 39 individuals and 664,522 and 1,106,859 loci, respectively.

To better understand F_ST_ distribution across the genome between ancient and modern samples and the drivers of temporal genetic diversity changes, we assessed F_ST_ outliers in OutFLANK v.02(118) using the unpruned filtered modern and ancient dataset of 90 individuals and 30,264 SNPs. OutFLANK was run in R using suggested settings (LeftTrimFraction = 0.05, RightTrimFraction = 0.05) and a false discovery rate (FDR) of ≤5%(118). OutFLANK accounts for sampling error and nonindependence between sample groups provided a priori while inferring the F_ST_ distribution of loci and fitting a χ2 model to the centre of the distribution.

## Data, Materials, and Software Availability

Prior to the final round of peer review, the raw genome sequence data reported in this article will be deposited in the European Nucleotide Archive. The genomic reference used to analyse the data is available at NCBI BioProject: PRJNA408269. All meta data are included in the article and/or supporting information.

## Author Contributions

AJA, FT, EC and AC designed the study. AJA, EFE, BS, FT, EM, GZ, VO, VA, GC, GP, SVN and PP collected samples for analysis. AJA, EFE, LA, OK, FP, FF and EC conducted the laboratory work. AJA, KP, LA, EFE and OK analysed the data. AJA, EFE, KP, BS, ADT, and FT wrote the paper. All authors reviewed the paper.

## Acknowledgements

This work is a contribution to the https://tunaarchaeology.org project within the framework of the MSCA SeaChanges ITN, which was funded by EU Horizon 2020 (Grant Number: 813383). We are grateful to Sara De Fanti and Anisa Ribani for laboratory support, and to Paolo Abondio and Shripathi Bhat for bioinformatics advice. The authors would like to thank the NOAA Restore Project which aided the collection of western Atlantic samples and the Institute of Marine Research (Norway) for access to modern Norwegian samples.

## Supporting Information

### Supporting text

#### Sample Details

Table 1 - external file Supplementary Table S1

##### Modern samples

Young-of-the-year Mediterranean samples were randomly selected from those caught in spawning areas as part of a separate study(26).

Gulf of Mexico larval samples were collected as part of the NOAA Restore project (https://restoreactscienceprogram.noaa.gov/projects/bluefin-tuna-larvae). Larvae were randomly selected for sequencing from three sampling surveys conducted in 2014, 2017 and 2018 from the north and western shelf of the Gulf of Mexico. Multiple years and locations were sampled to obtain maximum genomic variability among few final samples analysed (n=10). Full details of sampling are found in Table S2.

**Table S2.** Sampling details of the Gulf of Mexico (GoM) 2014-2018 samples sequenced herein, collected across multiple years and locations as part of the NOAA Restore project.

|  |  |  |  |  |  |  |  |  |  |  |
| --- | --- | --- | --- | --- | --- | --- | --- | --- | --- | --- |
| *ID* | *Station* | *Ship* | *Gear* | *Sample_No* | *SL_mm*<br>*etoh* | *developmental stage* | *depth collected, m* | *Longitude* | *Latitude* | *Date* |
| N1704 0108 | 4 | NOAA SHIP NANCY FOSTER | 90 cm quadrangular bongo, 505 mesh | D03760 | 7.19 | postflexion | 0-25 | - 87.77 73 | 26.1 198 | 10-May -17 |
| N1704 0501 | 21 | NOAA SHIP NANCY FOSTER | 90 cm quadrangular bongo, 505 mesh | D03791 | 5.5 | flexion | 0-25 | - 88.13 28 | 25.8 438 | 12-May -17 |
| N1802 1808 | 91 | NOAA SHIP NANCY FOSTER | 90 cm quadrangular bongo, 505 mesh | D04664 | not measured | larvae | 0-25 | - 87.24 97 | 28.3 327 | 15-May -18 |
| N1802 1809 | 91 | NOAA SHIP NANCY FOSTER | 90 cm quadrangular bongo, 505 mesh | D04664 | not measured | larvae | 0-25 | - 87.24 97 | 28.3 327 | 15-May -18 |
| N1802 1810 | 91 | NOAA SHIP NANCY FOSTER | 90 cm quadrangular bongo, 505 mesh | D04664 | not measured | larvae | 0-25 | - 87.24 97 | 28.3 327 | 15-May -18 |
| W1405 0007 | 23 | F.G. Walton Smith (UNOLS Ship) | 1x2m rectangular net, 505 mesh | 47654 | 4.28 | larvae | 0-10 | - 93.58 2 | 26.5 346 | 7-May -14 |
| W1405 0014 | 23 | F.G. Walton Smith (UNOLS Ship) | 1x2m rectangular net, 505 mesh | 47654 | 4.32 | larvae | 0-10 | - 93.58 2 | 26.5 346 | 7-May -14 |
| W1405 0029 | 27 | F.G. Walton Smith (UNOLS Ship) | 1x2m rectangular net, 505 mesh | 47659 | 5.05 | larvae | 0-10 | - 93.00 33 | 27.0 388 | 8-May -14 |
| W1405 0096 | 69 | F.G. Walton Smith (UNOLS Ship) | 1x2m rectangular net, 505 mesh | 47701 | 6.68 | larvae | 0-10 | - 87.76 1 | 27.9 963 | 20-May -14 |
| W1405<br>0097 | 69 | F.G. Walton<br>Smith (UNOLS<br>Ship) | 1x2m rectangular<br>net, 505 mesh | 47701 | 5.94 | larvae | 0-10 | -<br>87.76<br>1 | 27.9<br>963 | 20-May<br>-14 |

A total of 10 bluefin tuna were analysed from catches off Ålesund in 2018. These were adult individuals ca. 2 m in fork length (FL) and were analysed as part of a separate study, for full details see (84).

##### Ancient samples

Archival and archaeological sample details are listed below along with body size estimates calculated using the online tool https://tunaarchaeology.org/lengthestimations/ as detailed in the publication (119).

##### 1925 & 1941 tuna trap archived samples

We analysed specimens collected from two locations in the early 20^th^ century by the ecologist Massimo Sella (49). All specimens consist of vertebrae that were air-dried by the collator after capture and processing at tuna traps (Tonnare). A total of 8 vertebrae specimens were obtained from BFT captured in the tonnara at Zilten, Libya (Ionian Sea) in 1925, estimated to represent BFT between 158-204 cm FL, average 182 cm FL. Two large (275 and 278 cm FL) specimens were also sampled from tuna traps in the Bosporus, Istanbul, Turkey in 1941.

###### 1750-1800 Marseille Harbour, Marseille, France

A total of seven bluefin tuna bones (6 opercula and 1 vertebra) were obtained from the excavation Marseille-22 rue Jean François Leca, of the ancient port of Marsille, France, which was dated to between the late 18th and early 19th century (120). An approximate date of 1750-1850 CE is shown for this sample group in analyses. FL estimates were not made for these individuals as the vertebra selected was fragmented and could not be assigned to rank or accurately measured. Specimens represented large ca. 2 m sized adult bluefin tuna.

###### 1600-1800 Pedras de Fogu, Sassari, Italy

Ten vertebrae samples were obtained from the archaeological site of ‘Pedras de Fogu’ (Sassari, Sardinia, Italy). A tuna trap (tonnara) operated at this location from the end of the 16^th^ to the end of the 18^th^ century where BFT vertebrae have been recovered in a midden at the back of the beach after they were revealed by coastal erosion (121). These specimens were estimated to range from 115-231 FL, average 178 cm FL.

###### 900-1200 Sicily, Italy

A total of 3 specimens (2 vertebrae and 1 cranial element) were obtained from the archaeological site of ‘Mazara del Vallo’ situated in the town (southwestern Sicily, Italy). Samples were recovered from urban 10-13^th^ century layers, each dated by context as detailed in (122), and identified as different individuals according to their range of sizes. FL estimates were not made for these individuals as the vertebrae selected were fragmented and could not be assigned to rank or accurately measured. Broadly, specimens represented small-large sized adult bluefin tuna. A total of 3 samples (2 vertebrae and 1 cranial element) were selected for analyses from urban 9-10^th^ century layers in two different excavations in settlements in the city of Palermo, Sicily; Sant’Antonino and Corso dei Mille. The layers were dated by context as detailed in (122). Samples were estimated to represent individuals ranging from 101-185 cm FL, average 130 cm FL, believed to have been caught locally.

###### 800-1200 Yenikapi, Istanbul, Turkey

Ten vertebrae specimens were selected for analyses from an excavation at a Byzantine era site in the Yenikapi neighbourhood of Istanbul, Turkey. The Port of Theodosius operated at this site from 4-11^th^ century before being filled in the 15^th^ century (123). The 800-1200 CE origin of the samples is proposed from carbon dating achieved in a separate study(15). It is unknown whether specimens were fished locally or transported to the city of Constantinople, which was a major trading hub throughout the Byzantine period.

###### 3000 BCE Jortveit, Norway

A total of two samples were used from Jortveit, southern Norway. Accurate FL estimates could not be made for these individuals as the vertebrae were fragmented and could not be assigned to rank or measured accordingly. Nonetheless, the bones were of considerable size, likely representing specimens ca. 2 m sized adult BFT. These individuals were carbon dated to ca. 3000 BCE. For full details see (84)

### Supplementary Figures

#### Mapping and sample quality

**Figure S1.**
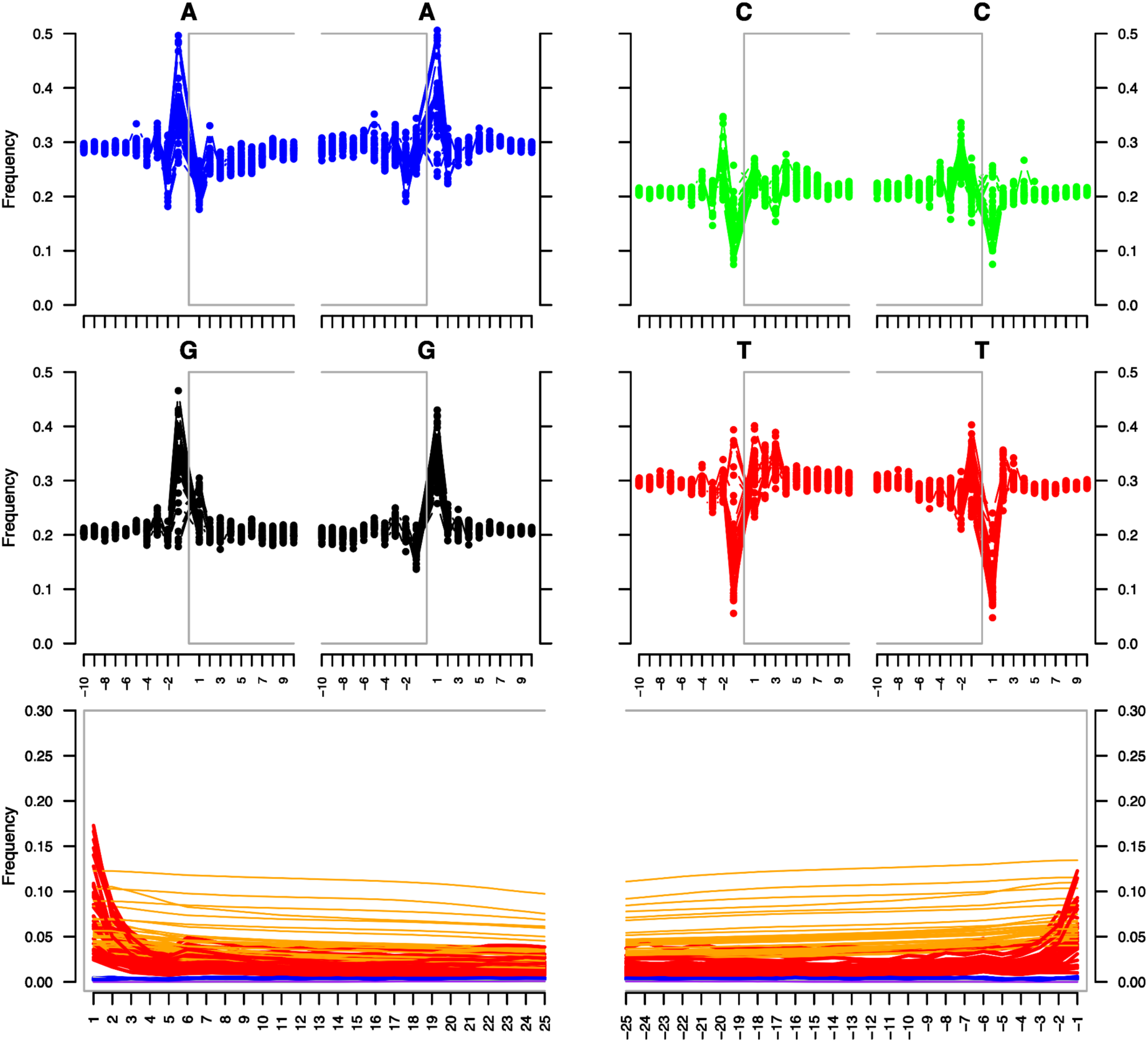
MapDamage fragment length plots for ancient Atlantic bluefin tuna (*Thunnus thynnus*) samples (n=45). X-axes represent base pair distance from terminal ends of sequence reads.

#### Demographic analyses

**Figure S2.**
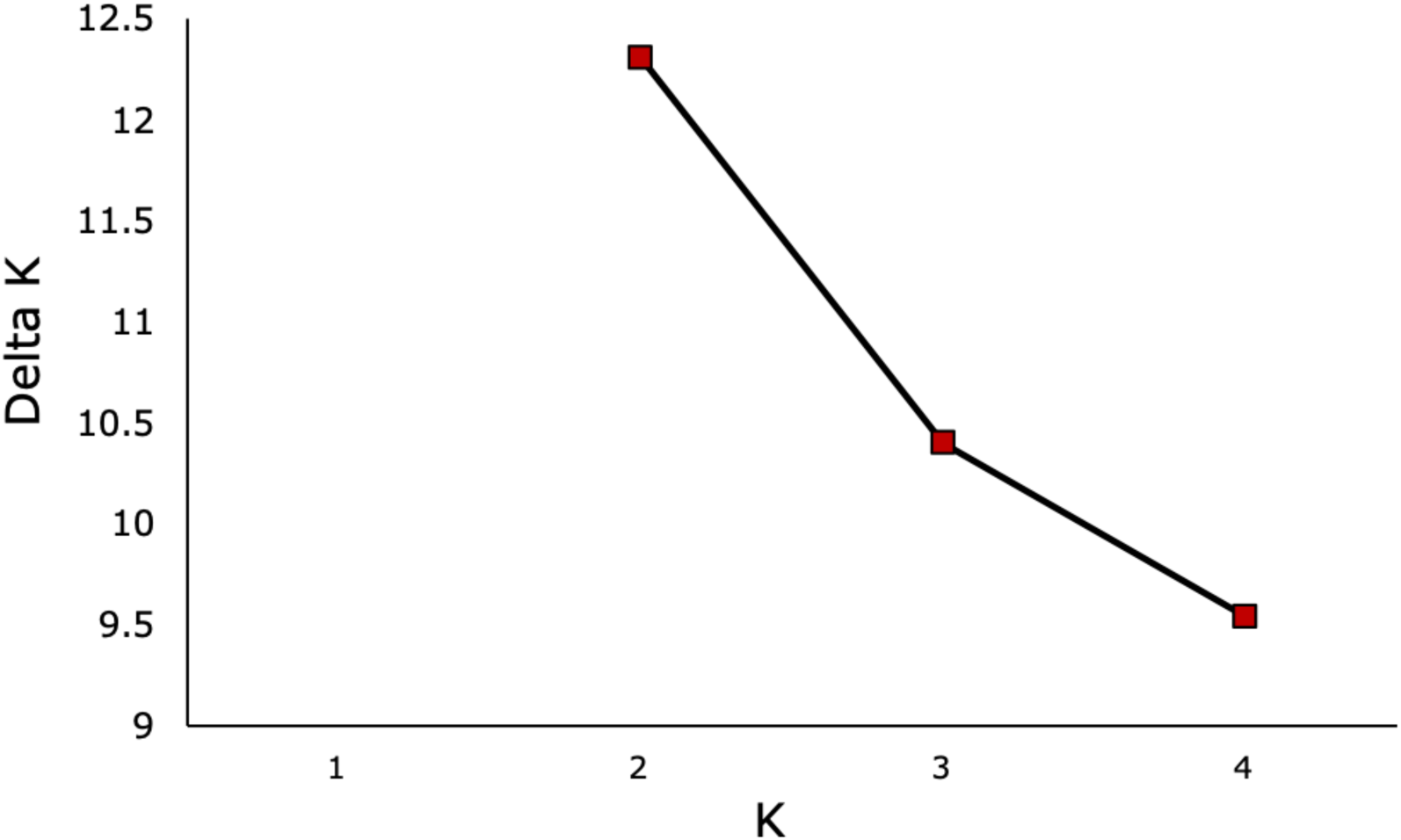
Delta K of 2 was selected for the optimal number of Atlantic bluefin tuna (*Thunnus thynnus*) populations using the modern spawning site dataset (n=39).

**Figure S3.**
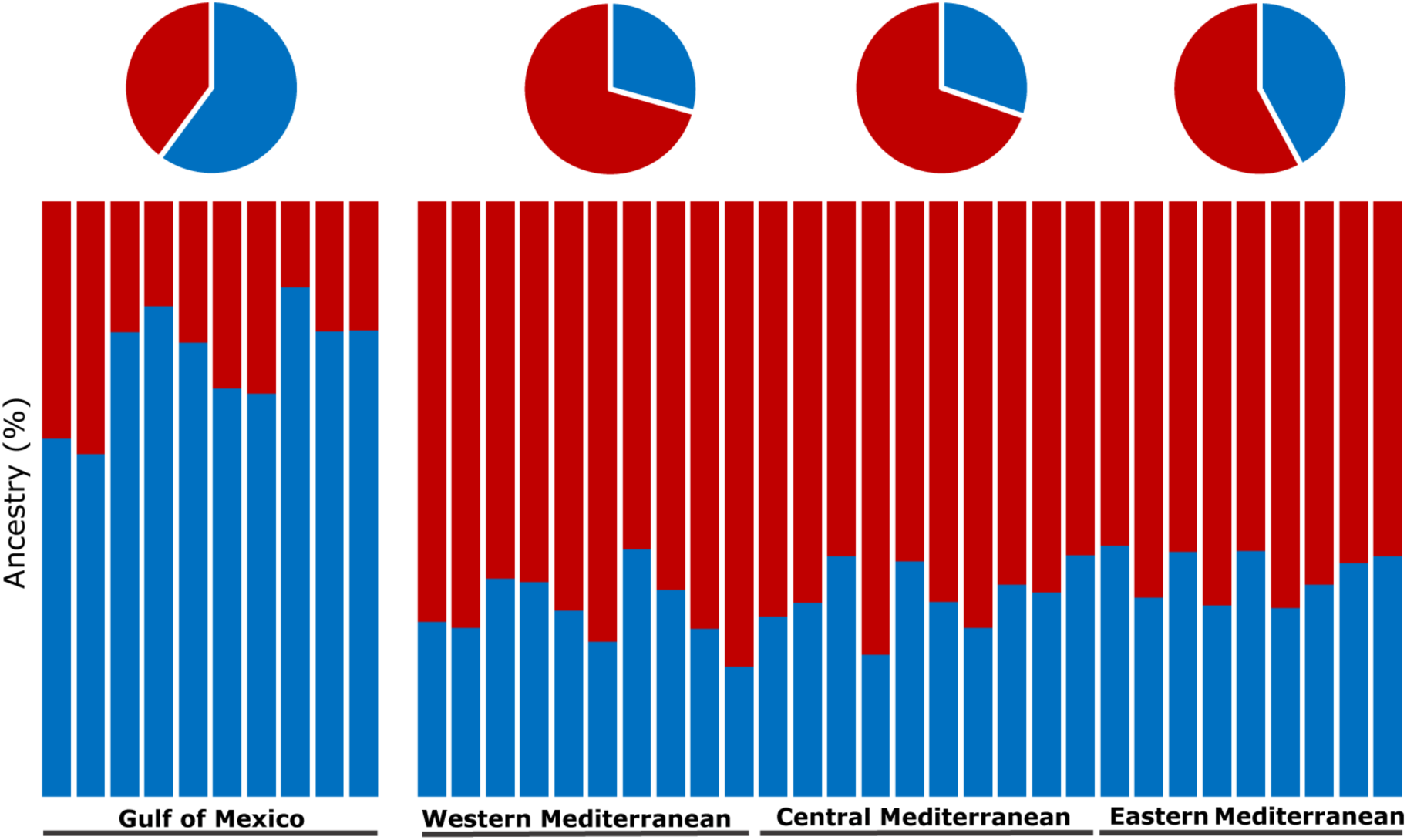
STRUCTURE bar plots and pie charts for the modern Atlantic bluefin tuna (*Thunnus thynnus*) (n=39) spawning site pruned dataset at 386,963 SNPs. Illustrating that at a greater number of loci not using F_ST_ site-filtering methods, the Eastern Mediterranean has a slightly greater quantity of Gulf of Mexico ancestry than Western and Central Mediterranean bluefin tuna, as discovered using the modern and ancient combined dataset in the main paper (Figure 1).

**Figure S4.**
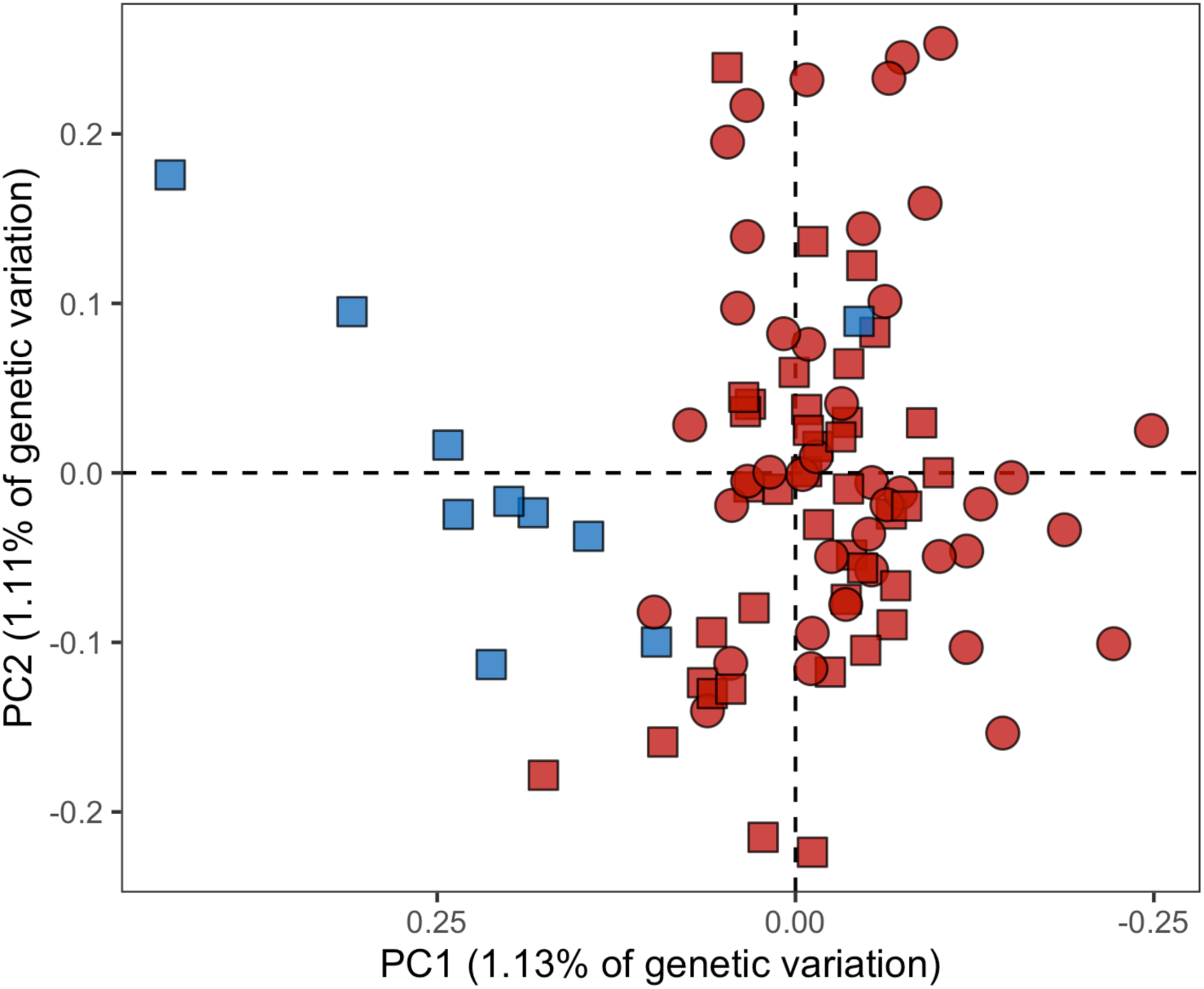
Principal components analysis (PCA) of the modern and ancient dataset at 27,882 SNPs which have not been filtered using the F_ST_ site-filtering approach. Illustrating that without our approach, PCAs cannot clearly separate the two populations of Atlantic bluefin tuna (*Thunnus thynnus*).

**Figure S5.**
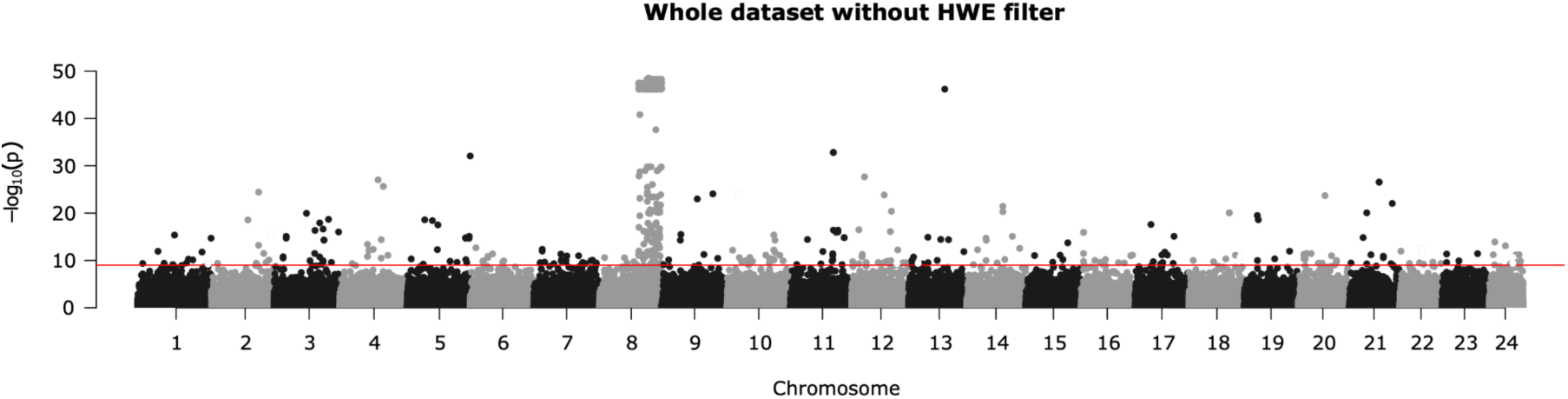
Manhattan plot of pcadapt outlier detection results for Atlantic bluefin tuna (*Thunnus thynnus*) (n=39) using the modern spawning site dataset to show introgression signals on Chromosome 8 when including loci that deviate from Hardy Weinberg Equilibrium.

**Figure S6.**
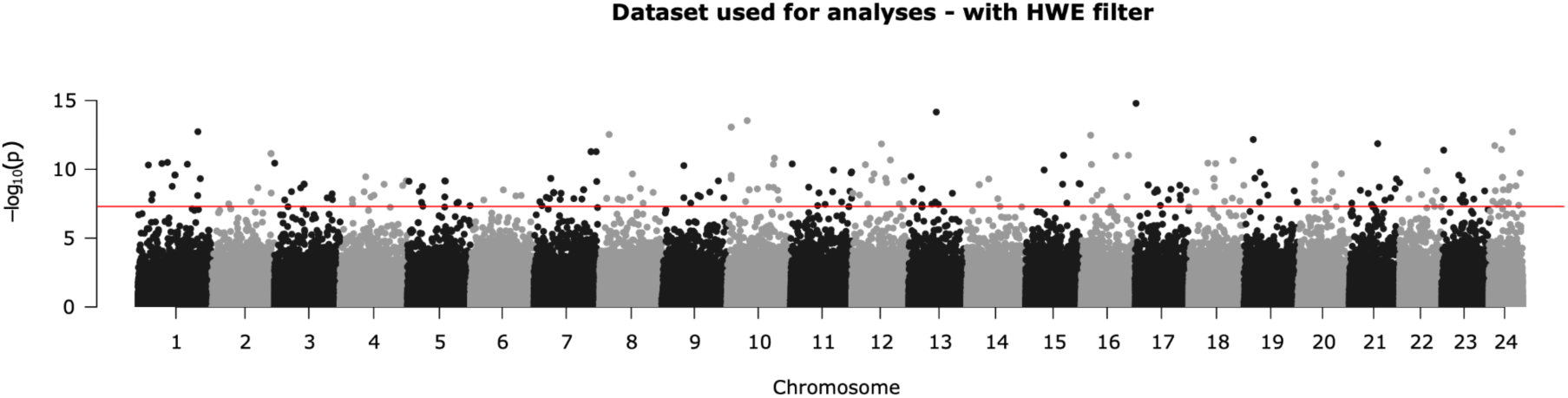
Manhattan plot of pcadapt outlier detection results for Atlantic bluefin tuna (*Thunnus thynnus*) (n=39) using the modern spawning site dataset to show the removal of introgression signals on Chromosome 8 when removing loci that deviate from Hardy Weinberg Equilibrium. See Methods. This approach was used for all demographic analyses.

**Figure S7.**
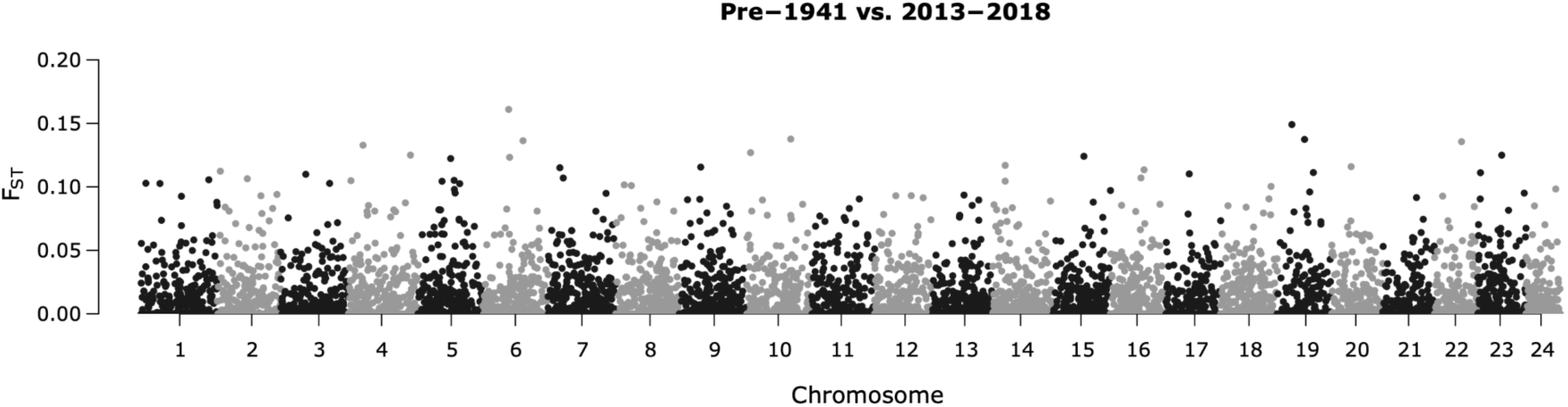
Manhattan plot of Pcadapt outlier detection results for Atlantic bluefin tuna (*Thunnus thynnus*) (n=90) using the combined modern and ancient dataset to show the distribution of F_ST_ temporal differences across the genome being relatively uniform and low F_ST_.

**Figure S8.**
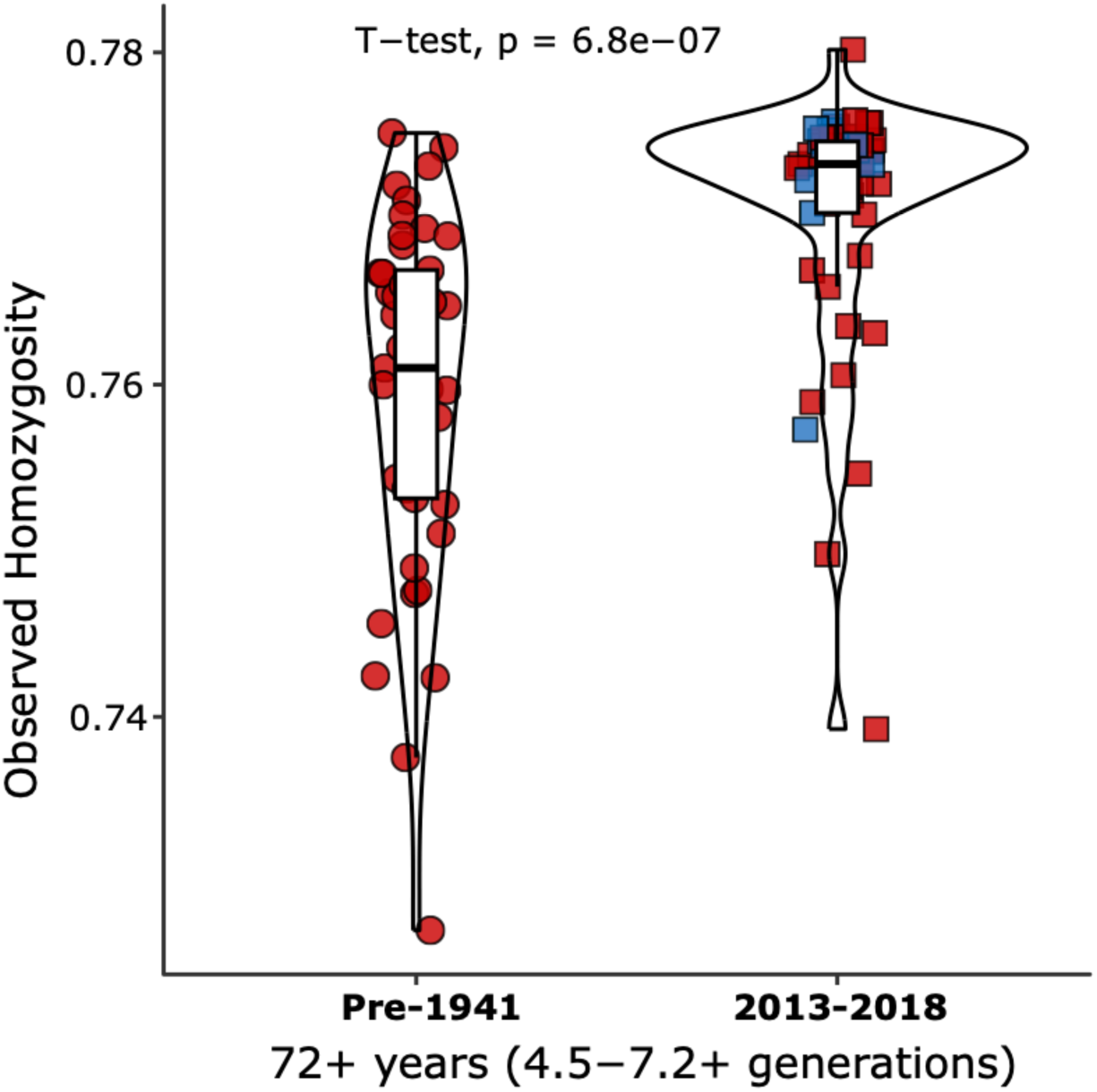
Violin plots show observed homozygosity (fraction of sites) differences when F_ST_ site-filtering is not applied to the dataset and 10% missing data is allowed at 2,890,241 SNPs between pooled ancient (Pre-1941, n=41) and modern (2013-2018, n=49), coloured by population (blue: western stock, red: eastern stock) and with shapes depicting time period (circle: ancient, square: modern).

**Figure S9.**
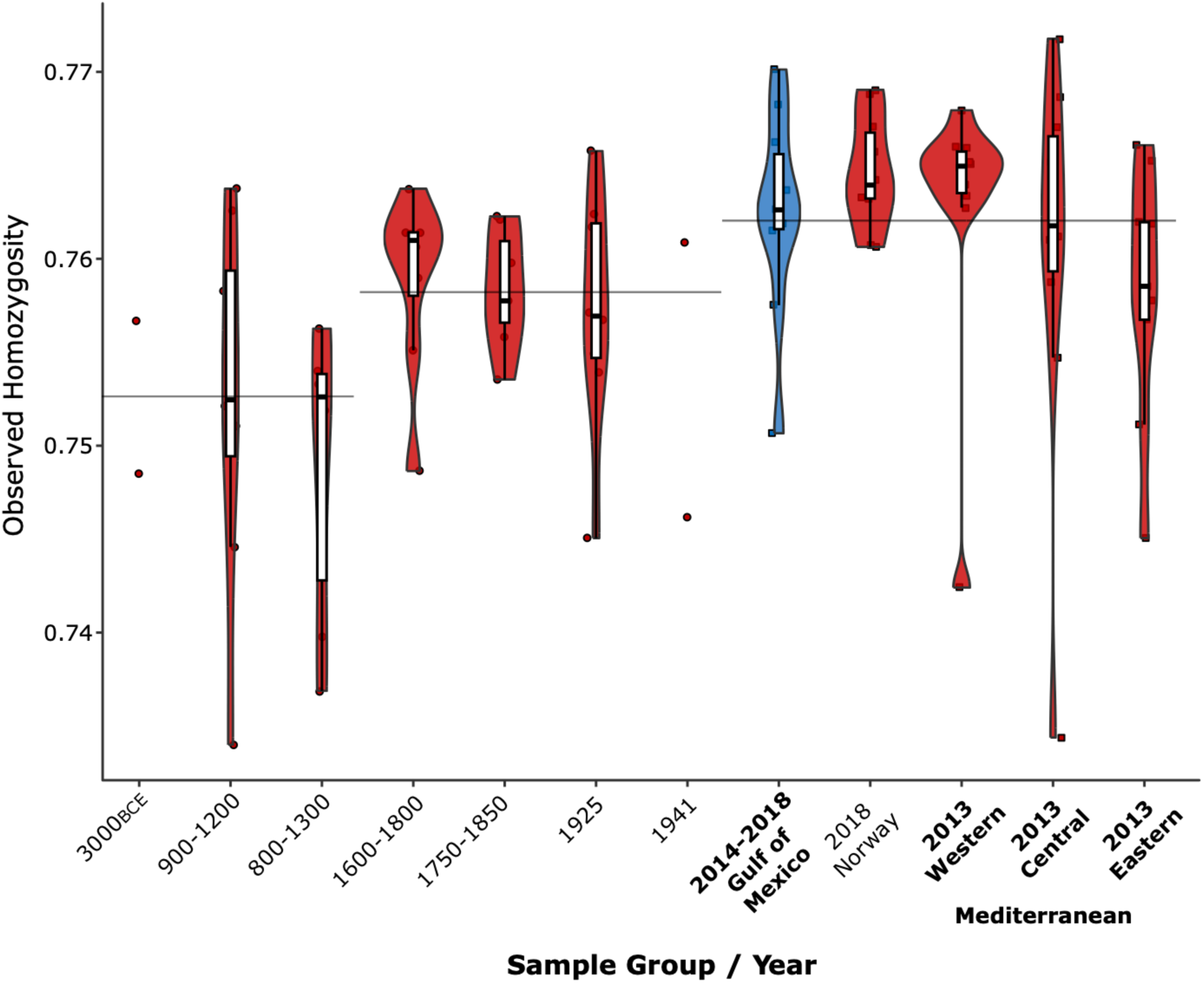
Violin plots show step-wise loss of observed homozygosity (fraction of sites) over time at 12,838 SNPs between grouped ancient and modern Atlantic bluefin tuna (*Thunnus thynnus*) samples (n=90), coloured by stock (blue: western stock, red: eastern stock) and with shapes depicting time period (circle: ancient, square: modern). Boxplots are illustrated for groups >2 sample size, with group means, 25^th^ and 75^th^ percentile as outer edges and 95^th^ percentiles (black whiskers). Horizontal black lines between groups of time point samples (3000BCE to 900-1300; 1600-1800 to 1941; modern 2013-2018 samples) show averages.

**Figure S10.**
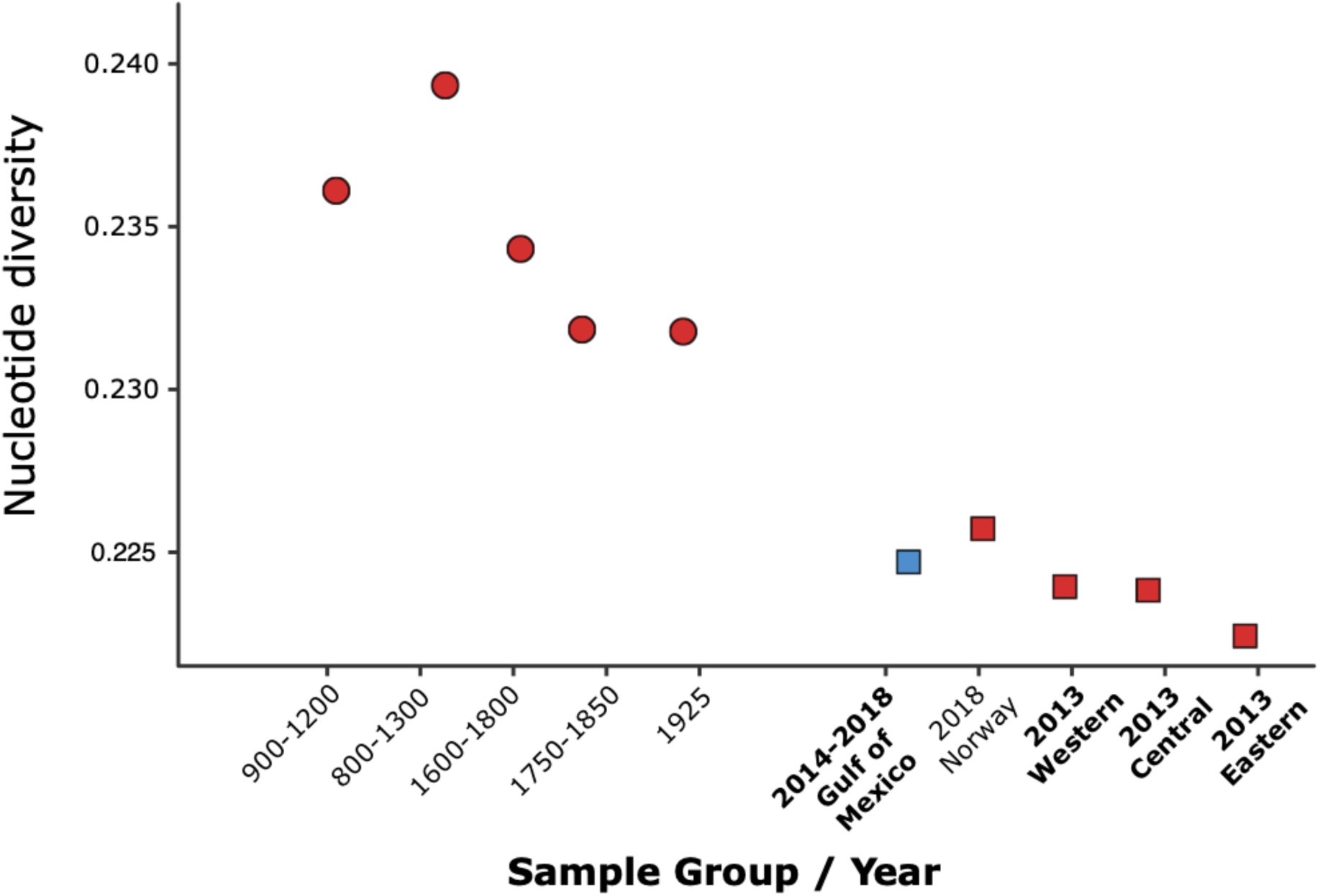
Nucleotide diversity estimates for each sample group show the trajectory of pre-industrial decline between ancient and modern Atlantic bluefin tuna (*Thunnus thynnus*), calculated including missing data at 2,890,241 loci, estimating nucleotide diversity average of 50Kb windows and not showing groups with sample size <2. Groups coloured by stock (blue: western stock, red: eastern stock).

**Figure S11.**
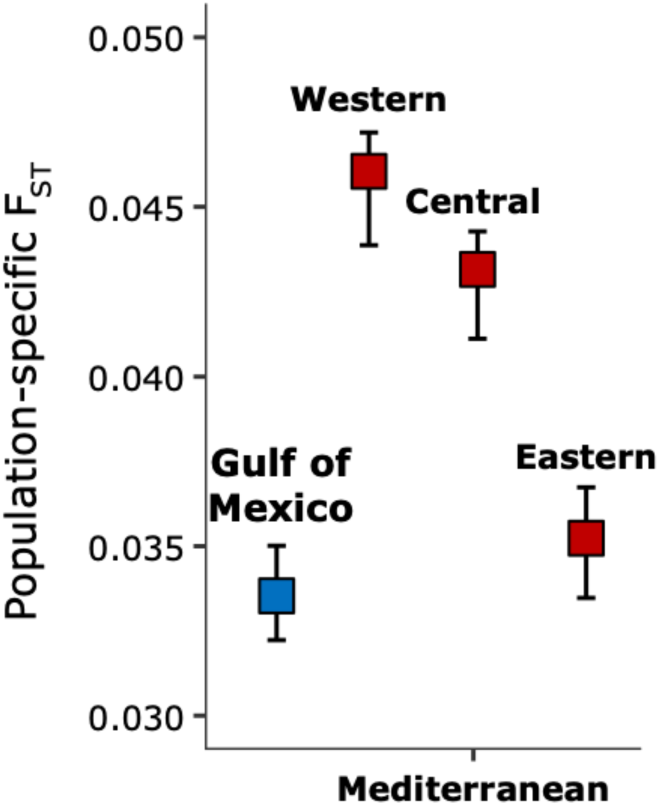
Error bars show population-specific F_ST_ estimates for each modern spawning site sample at 386,963 SNPs, with 95% confidence intervals (CIs, black whiskers).

**Figure S12.**
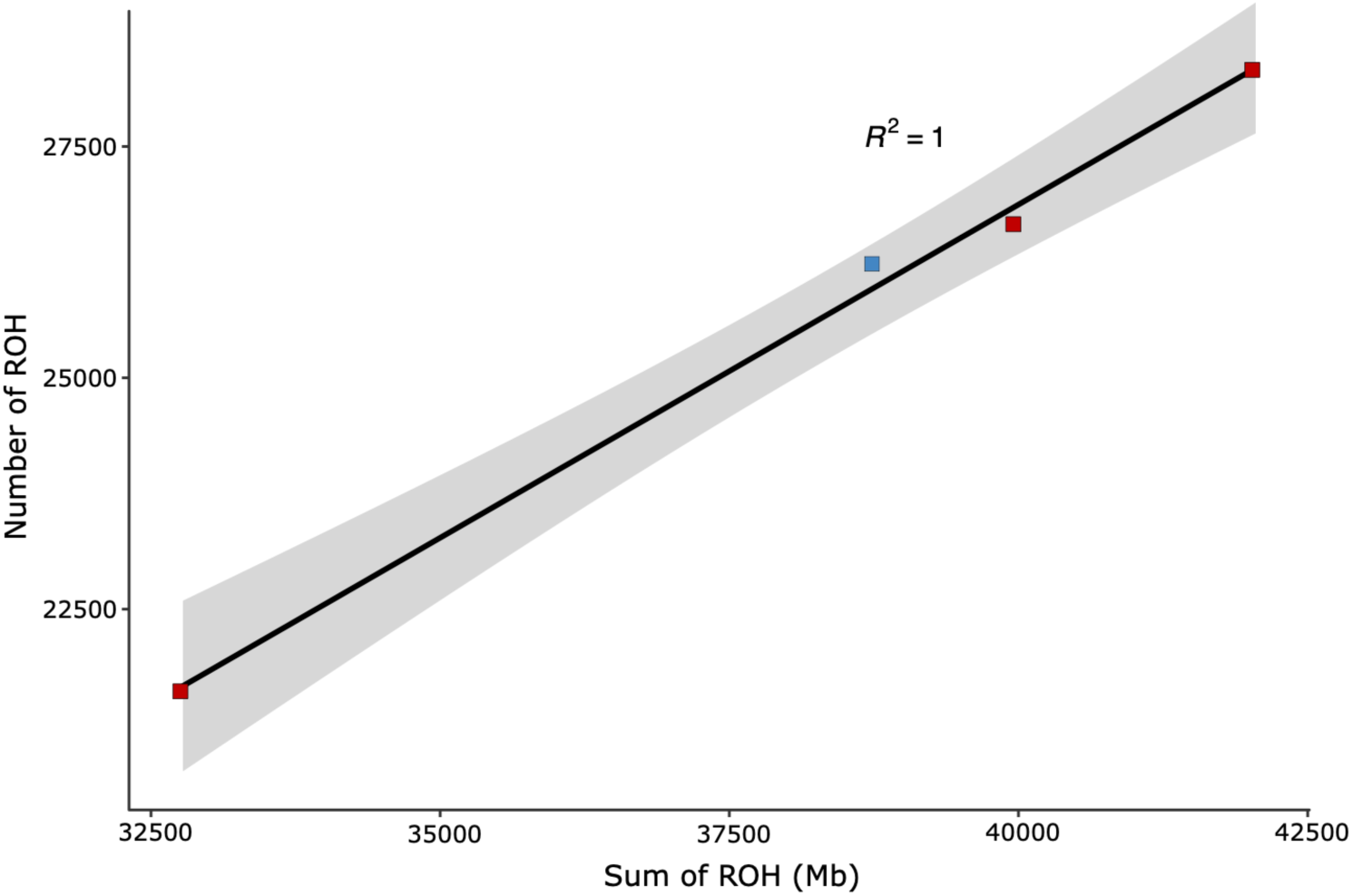
Scatter plot of the number of runs of homozygosity (NROH) vs the sum of runs of homozygosity (SROH) with geom_smooth fit line (black, grey shading as 95% confidence interval) for modern Atlantic bluefin tuna (*Thunnus thynnus*) samples analysed (n=39). Correlation between NROH and SROH) was strong as shown by R2=1.

**Figure S13.**
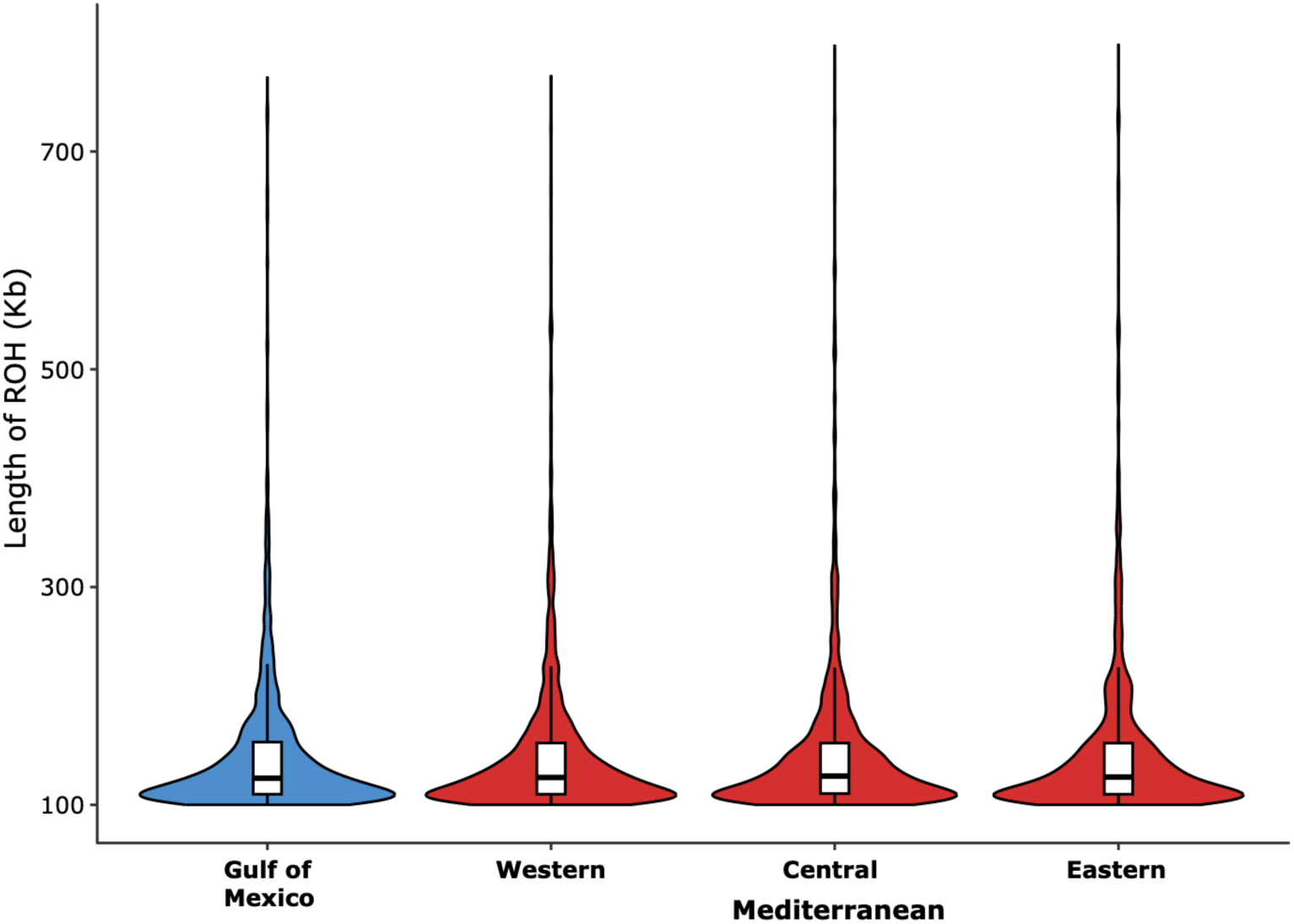
Violin plots no difference between Atlantic bluefin tuna (*Thunnus thynnus*) modern spawning site samples (n=39) in lengths of runs of homozygosity (ROH), coloured by stock (blue: western stock, red: eastern stock). Boxplots are illustrated with group means, 25^th^ and 75^th^ percentile as outer edges and 95^th^ percentiles (black whiskers).

**Figure S14.**
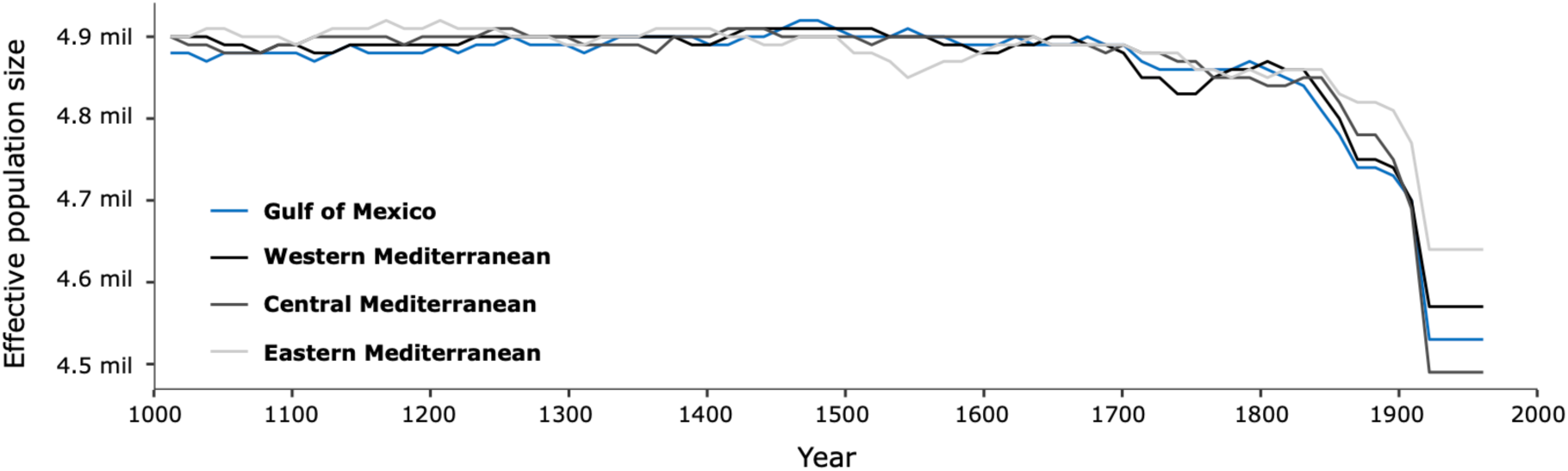
GONE estimates of historical effective population size (N_e_) for grouped Atlantic bluefin tuna (*Thunnus thynnus*) samples (n=39) confirm similar trajectories between Mediterranean sites as when pooled (main paper, Figure 2). Mediterranean spawning sites (grey scale) and Gulf of Mexico (blue), representing the previous 100 generations from 1000 CE to 1964, using the most likely generation length of 13 years.

**Figure S15.**
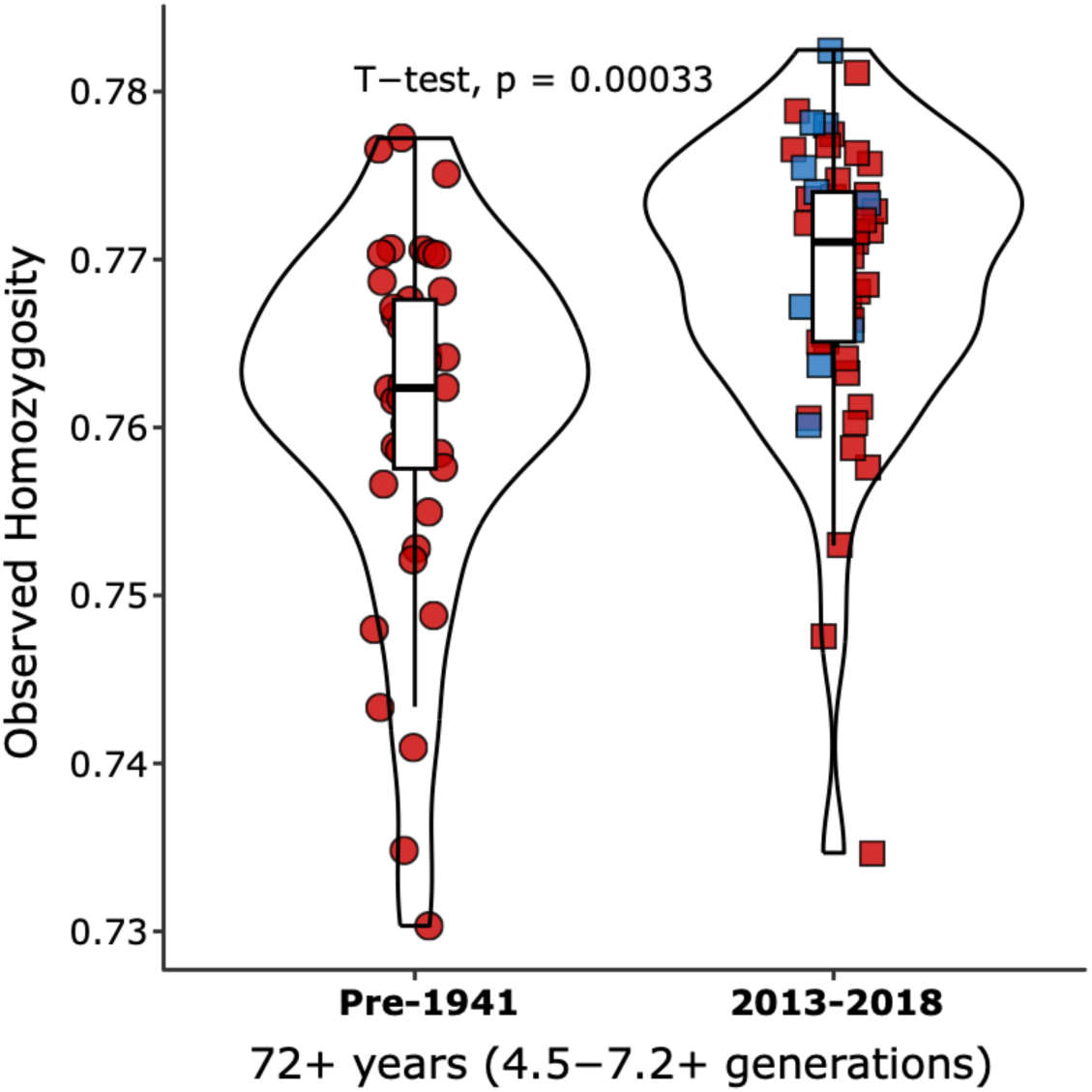
Violin plots show observed homozygosity (fraction of sites) differences when transition sites potentially subject to ancient DNA damage are excluded from the dataset at 5822 SNPs between pooled ancient (Pre-1941, n=41) and modern (2013-2018, n=49), coloured by population (blue: western stock, red: eastern stock) and with shapes depicting time period (circle: ancient, square: modern).

**Figure S16.**
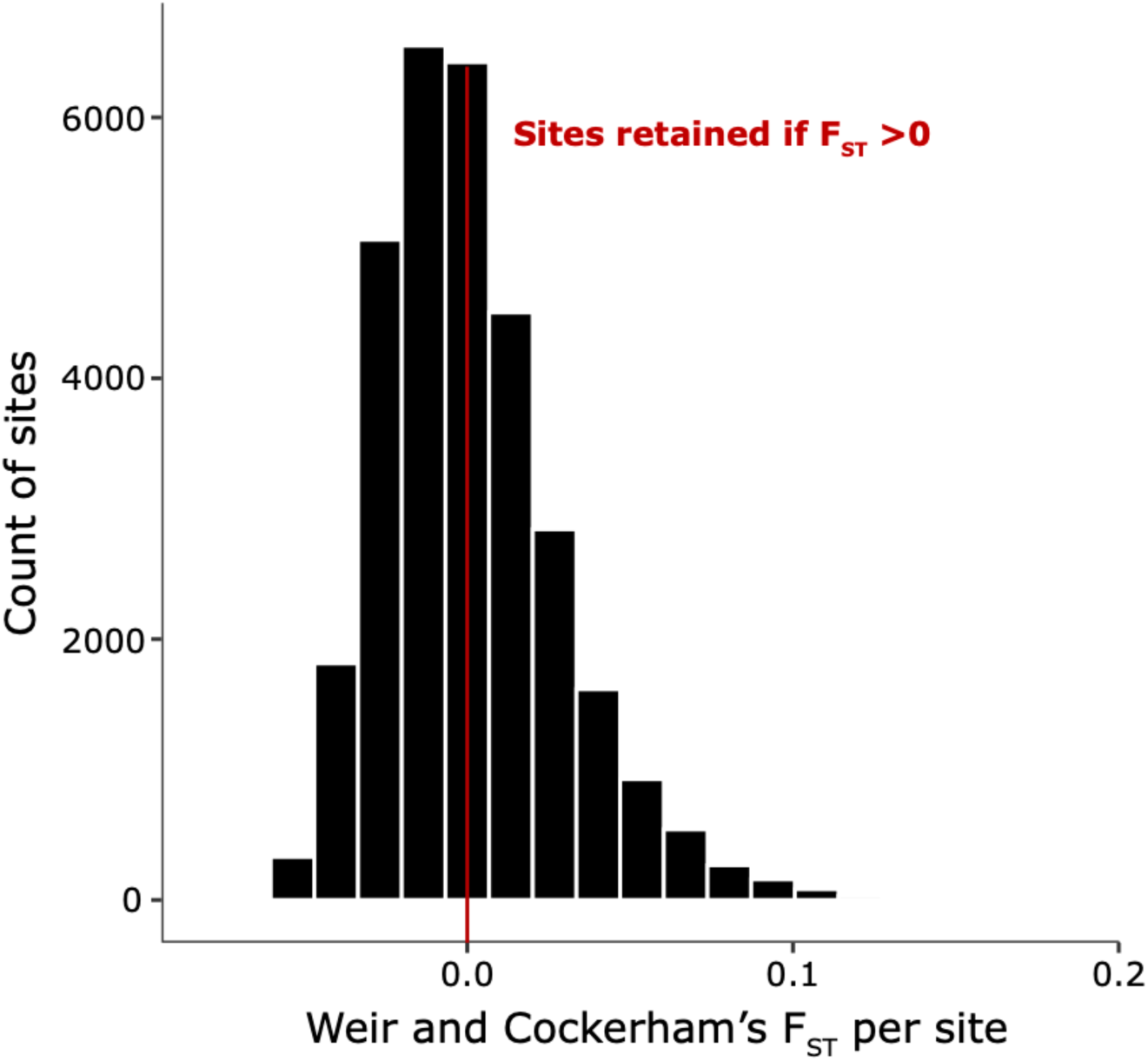
Histogram of per-site F_ST_ values calculated using Weir and Cockerhams F_ST_ in VCFTools. Illustrated is the F_ST_ distribution of the modern and ancient dataset at 30,264 SNPs showing cut off used for filtering (0 F_ST_).

